# The ESCRT-III machinery participates in the production of extracellular vesicles and protein export during *Plasmodium falciparum* infection

**DOI:** 10.1101/2020.11.24.395756

**Authors:** Yunuen Avalos-Padilla, Vasil N. Georgiev, Elena Lantero, Silvia Pujals, René Verhoef, Livia N. Borgheti-Cardoso, Lorenzo Albertazzi, Rumiana Dimova, Xavier Fernàndez-Busquets

## Abstract

Infection with *Plasmodium falciparum* enhances extracellular vesicles (EVs) production in parasitized red blood cells (pRBC), an important mechanism for parasite-to-parasite communication during the asexual intraerythrocytic life cycle. The <u>e</u>ndosomal <u>s</u>orting <u>c</u>omplex <u>r</u>equired for <u>t</u>ransport (ESCRT), and in particular the ESCRT-III sub-complex, participates in the formation of EVs in higher eukaryotes. However, RBCs have lost the majority of their organelles through the maturation process, including an important reduction in their vesicular network. Therefore, the mechanism of EV production in *P. falciparum-*infected RBCs remains to be elucidated. Here we demonstrate that *P. falciparum* possesses a functional ESCRT-III machinery that is activated by an alternative recruitment pathway involving the action of PfBro1 and PfVps32/PfVps60 proteins. Additionally, multivesicular bodies formation and membrane shedding, both reported mechanisms of EVs production, were reconstituted in the membrane model of giant unilamellar vesicles using the purified recombinant proteins. Moreover, the presence of PfVps32, PfVps60 and PfBro1 in EVs purified from a pRBC culture was confirmed by super-resolution microscopy. In accordance, disruption of the *Pfvps60* gene led to a reduction in the number of the produced EVs in the KO strain when compared with the parental 3D7 strain. Overall, our results increase the knowledge on the underlying molecular mechanisms during malaria pathogenesis and demonstrate that ESCRT-III *P. falciparum* proteins participate in EVs production.

## Introduction

*Plasmodium spp* is the parasite responsible for malaria, a disease that, despite the efforts done to control it, still represents a health problem worldwide particularly in developing countries [1]. During *Plasmodium* infection, an elevated number of extracellular vesicles (EVs) from numerous cellular sources are circulating in the plasma [2], the amount of which correlates with the severity of the disease [2–5].

Despite of its high impact in the development of the pathology, the precise mechanism of EV formation in the infected red blood cells (RBCs) remains to be elucidated. One of the yet unsolved enigmas of malaria pathophysiology is how mature RBCs are able to release high amounts of EVs after *Plasmodium* infection, since they are biochemically simple compared to other eukaryotic cells and lack a normal vesicular network. It has been suggested that *Plasmodium* uses its own protein network to establish a vesicular trafficking for the export of an arsenal of virulence factors through which contributes to the establishment of the parasite into the host cells [6].

In higher eukaryotes, EVs are generated and transported to their final destination by the endomembrane system [7]. Trafficking within the endomembrane system is crucial for the functional communication between different compartments in eukaryotic cells [8]. Depending on their origin and size, EVs can be classified into two major classes: exosomes and microvesicles. Exosomes refer to endosome-derived vesicles with a diameter typically of 30-50 nm that are generated following the fusion of multivesicular bodies (MVBs) with the plasma membrane. On the other hand, microvesicles are plasma membrane-derived vesicles which result from direct membrane shedding and exhibit a size from 100 nm up to 1 μm [9].

The MVBs are shaped after the formation of intraluminal vesicles (ILVs) in early endosomes [10]. The genesis of ILVs relies on the sequential action of the <u>e</u>ndosomal <u>s</u>orting <u>c</u>omplex <u>r</u>equired for <u>t</u>ransport (ESCRT), which consists of four protein complexes termed ESCRT-0, ESCRT-I, ESCRT-II, ESCRT-III and a set of accessory proteins [11, 12]. The best-described mechanism of ESCRT action begins with the recognition of mono-ubiquitinated proteins by ESCRT-0 [13], which then activates the recruitment of ESCRT-I [14] and ESCRT-II [15] that are responsible for membrane deformation into buds [16, 17]. Finally, the polymerization of ESCRT-III begins with the binding of Vps20 to the invaginated membrane, which recruits the rest of the ESCRT-III members to the bud neck and the nascent vesicle is closed [16, 18, 19]. The dissociation and recycling of the machinery depends on the participation of the Vps4 AAA-ATPase [20]. Among all the sub-complexes, ESCRT-III (composed by Vps20, Snf7/Vps32, Vps24 and Vps2) and its accessory proteins (Vps4, Vta1, Vps60, ALIX) are also involved in other important membrane-scission mechanisms, including virus budding, cytokinesis, nuclear envelope remodeling and exosome biogenesis among others (see review in [21]). All of these processes share the same topology where the nascent vesicle buds away from the cytosol, contrary to the topology observed in clathrin-coated vesicles [22].

The ESCRT machinery is highly conserved across the eukaryotic lineage; however, strictly intracellular protists, like *Plasmodium spp*, are devoid of ESCRT-0, -I and -II sub complexes [23]. In the case of *Plasmodium* and other organisms that lack the full ESCRT machinery, it is plausible that other proteins trigger ESCRT-III activation. In this regard, ALIX, a Bro1-domain protein, binds directly to Vps32 and triggers the formation of ESCRT-III polymers, leading to ILVs formation in humans [24]. Whether a similar mechanism, alternative to the canonical ESCRT-III pathway, exists in *Plasmodium* remains to be determined.

Previous *in silico* assays showed that *Plasmodium falciparum,* the deadliest human malaria parasite species, possesses at least two putative proteins from the ESCRT-III complex: Vps2 and Vps32/Snf7 [23, 25]. Additionally, the ATPase Vps4, an accessory protein of the ESCRT-III complex, was found in the cytoplasm of *P. falciparum* during the trophozoite blood stage [26]. Moreover, PfVps4 retained its function in MVBs formation when transfected into *Toxoplasma gondii* and COS cells, thus strongly suggesting the existence of a functional ESCRT machinery in *P. falciparum* that mediates the production of MVBs [26].

Since *P. falciparum* lacks upstream ESCRT complexes, here we have demonstrated the presence of a Bro1-domain protein (PfBro1) involved in an alternative recruitment pathway. In addition, the action of PfBro1 and two Snf7 homologues in membrane-buds formation was reconstituted using giant unilamellar vesicles (GUVs) as a model system [27] in which we have visualized the assembly sequence and the function of the proteins. Additionally, we have used a microinjection approach that allowed us to recreate the topology occurring in living cells and to study EVs formation using the purified recombinant proteins from the parasite. Moreover, we were able to detect the presence of all studied proteins in EVs secreted by *P. falciparum*-infected RBCs and inactivate one ESCRT-III related gene, which confirms its participation in EV biogenesis. Overall, our findings provide an important insight into protein export in *Plasmodium-*infected RBCs mediated by the parasite and describe a molecular target with antibody susceptibility that can be part of future vaccination or therapeutic strategies.

## Results

### *Plasmodium falciparum* possesses a Bro1 domain-containing protein

A previous *in silico* study of the *P. falciparum* genome revealed the presence of only six out of the 26 ESCRT-machinery proteins present in humans. The study showed that the genome of *P. falciparum* encodes four Snf7-domain containing proteins [23], a conserved feature in all ESCRT-III members [28]. Based on our *in silico* Basic Local Alignment Search Tool (BLAST) analysis, the four proteins were denoted as PfVps32, PfVps60, PfVps2 and PfVps46 (S1 Table).

The absence of ESCRT-I- and -II-associated genes and of a Vps20 homologue in the genome of *P. falciparum*, suggested the existence of an alternative recruitment pathway in the parasite. Hence, we explored the presence of a Bro1 domain-containing protein in *P. falciparum* that could bind directly to the Snf7 candidates and trigger the activation of the ESCRT-III system in this parasite, similarly to the process regulated by ALIX in humans.

An *in silico* search of the *P. falciparum* genome (http://www.plasmodb.org) showed that the parasite has a unique Bro1-containing homologue termed PF3D7_1224200 (hereafter referred to as PfBro1) with a 3175 bp open reading frame and carrying 4 introns. The open reading frame of PfBro1 encodes an 819 amino acid protein with a predicted molecular mass of 98,714 Da. Our further assays revealed that the amino acid sequence of full-length PfBro1 had an identity of 21.8% with ALIX, whereas the Bro1 domain in PfBro1 exhibited a 23.6% identity with its human homologue. Despite of this low amino acid conservation, we identified several conserved residues of the two charged polar clusters which, in several Bro1 homologues, stabilize the Bro1 domain [29]. These residues include R51, Y70 and E116 from the first cluster, and E187 and K246 from the second cluster (S1 Fig). Importantly, PfBro1 showed the conservation of the residue I144 (S1 Fig), which has been demonstrated to directly participate in the binding of Vps32 in *Saccharomyces cerevisiae* [29]. We then performed additional tertiary structure prediction assays, revealing that the full-length PfBro1 has a hypothetical hydrophobic tail in its C-terminal region (S2 Fig), which makes it a good candidate for the recruitment of ESCRT-III components at the level of the membrane.

### PfBro1 and PfVps32 are exported to the cytoplasm of the erythrocyte

In order to continue our characterization of the ESCRT-III machinery in *P. falciparum*, we focused on resolving the putative role of three proteins: (1) PfVps32, the most abundant protein in the ESCRT machinery; (2) PfVps60, whose human homologue, CHMP5, is able to bind directly to Brox [30], a Bro1-containing protein found in exosomes of human urine [31], which trigger their redistribution to membrane-enriched fractions [30]; and (3) PfBro1 as their potential recruiter and activator. Consequently, genes encoding the aforementioned proteins were synthesized and cloned into the appropriate vector to induce and purify the corresponding proteins (S3 Fig) which were used for the rest of the experiments.

Rabbit polyclonal antibodies against the purified recombinant proteins were generated and used to detect their presence in protein extracts obtained by detergent fractionation from *P. falciparum* cultures during the intraerythrocytic stage. PfVps32 was present in the saponin extracts containing RBC-cytosolic proteins, PfVps60 was found in the RIPA-fraction where most cytoskeletal components are present, and PfBro1 was found in the Triton X-100 extracts enriched in proteins from membranes and organelles (Fig 1a-c). Interestingly, while PfBro1 and PfVps32 migrated at the expected molecular weight (98 and 26 kDa, respectively), PfVps60 migrated at a higher molecular weight (46 kDa) than that calculated from its amino acid sequence (27 kDa) (Fig 1c, S3 Fig). This effect has been observed in other Snf7-containing proteins and is attributed to the high electric charge of the protein [32]. Western blot results indicated that PfVps32, PfVps60 and PfBro1 are expressed throughout the whole intraerythrocytic cycle (Fig 1a-d, 2a-c). PfVps60 and PfBro1 showed a similar expression pattern with a peak during the trophozoite stage, 32 h post invasion (hpi) (Fig 1d). On the other hand, the highest PfVps32 levels were observed at ring stages (0-16 hpi), with an important decrease towards the trophozoite stage (32-40 hpi) (Fig 1a, d).

**Fig 1.**
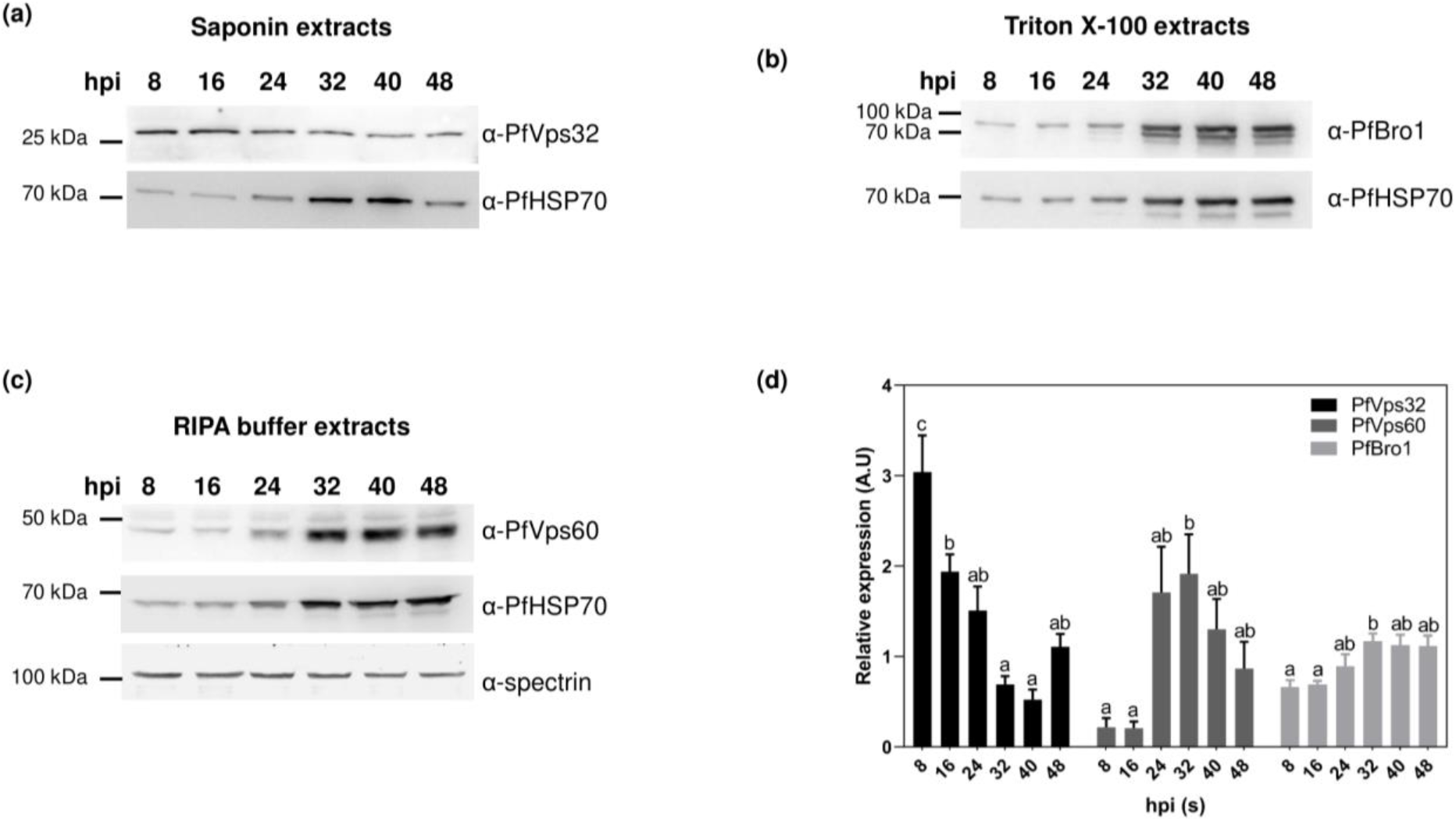
Expression of PfVps32, PfVps60 and PfBro1 during the *P. falciparum* intraerythrocytic cycle. Western blot analysis of *P. falciparum* in **(a)** saponin, **(b)** Triton X-100 or **(c)** RIPA buffer protein extracts at different hpi to monitor protein expression. **(d)** Intensities of the bands from PfVps32, PfVps60 and PfBro1 quantified by densitometry and normalized to those from PfHSP70, a ubiquitous *Plasmodium* protein present at all times tested. Data are means ± SE of 3 different replicates. Significant differences (*p* < 0.05) calculated using one-way ANOVA are denoted by different letters: shared letters represent no statistically significant difference.

**Fig 2.**
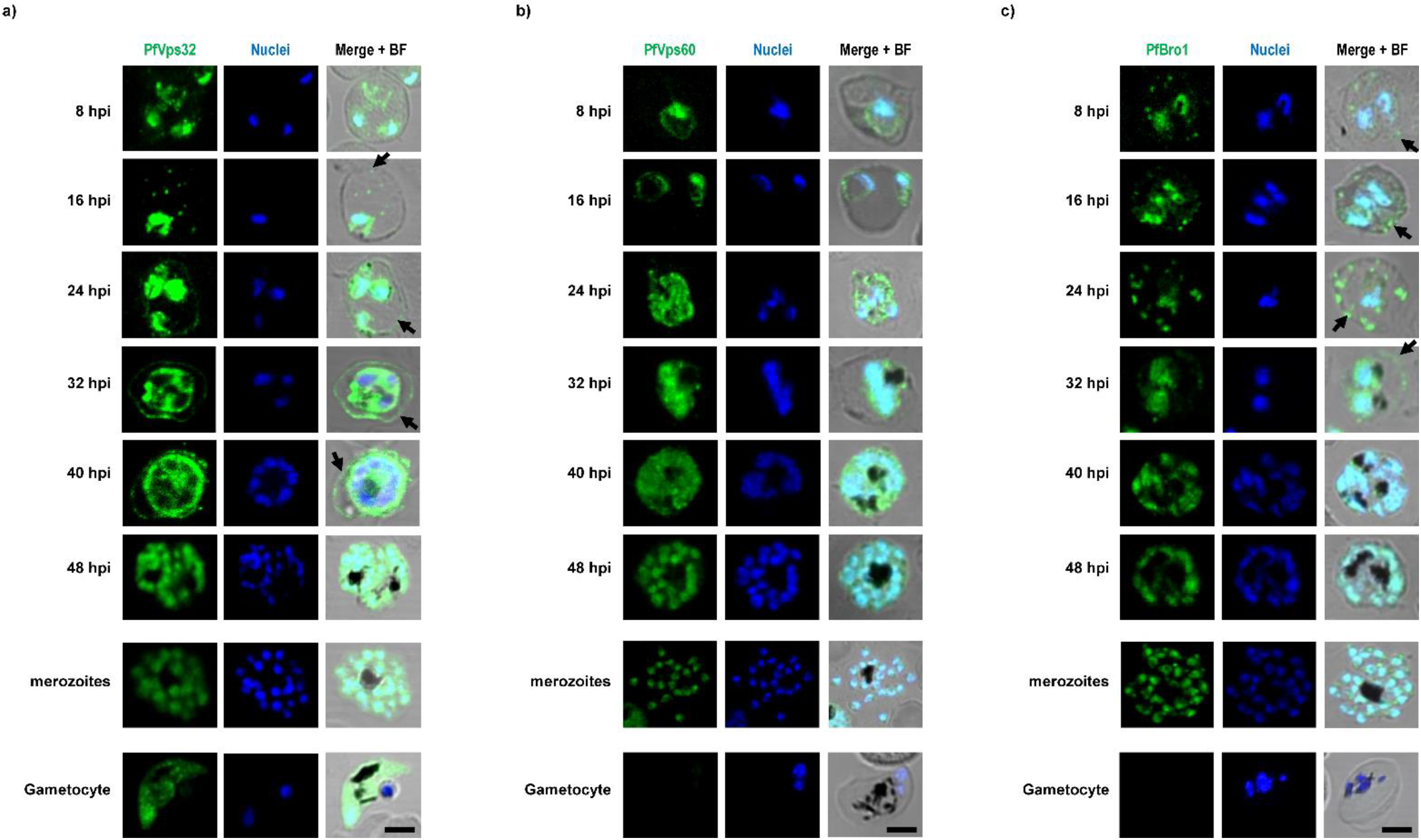
Subcellular localization of PfVps32, PfVps60 and PfBro1 during the *P. falciparum* intraerythrocytic cycle. Human erythrocytes infected with *P. falciparum* were fixed at different hpi and **(a)** PfVps32, **(b)** PfVps60 and **(c)** PfBro1 were detected by indirect immunofluorescence microscopy. For gametocyte imaging only one stage is showed (see S1 Text). Cell nuclei were visualized with Hoechst 33342 (blue). Fields were merged with bright field (BF) to assess localization. Scale bar: 5 μm.

Interestingly, antibodies against PfBro1 and PfVps60 detected more than one band, which suggested a proteolytic processing of the proteins or its association in a complex probably related with their function or degradation (S4 Fig).

Immunofluorescence assays showed that PfVps32, PfVps60 and PfBro1 were localized in the cytoplasm of the parasite (Fig 2). In the case of PfVps32, the protein was detected on the membrane of the infected RBC from 24 to 40 hpi (Fig 2a), indicating its export during these times. Moreover, stage I to stage IV gametocytes stained positive for PfVps32 whereas PfBro1 and PfVps60 were absent from these stages. Additionally, punctate structures stained with anti-PfBro1 and PfVps32 were observed in the cytoplasm of parasitized erythrocytes outside the parasitophorous vacuole (Fig 2c arrows). To examine whether these structures are exported to the parasitized RBC (pRBC) plasma membrane, lectins present in the RBC surface were labeled with wheat germ agglutinin (WGA). PfVps32 and PfBro1-labeled vesicles co-localized with the surface lectin (Fig 3, arrows).

**Fig 3.**
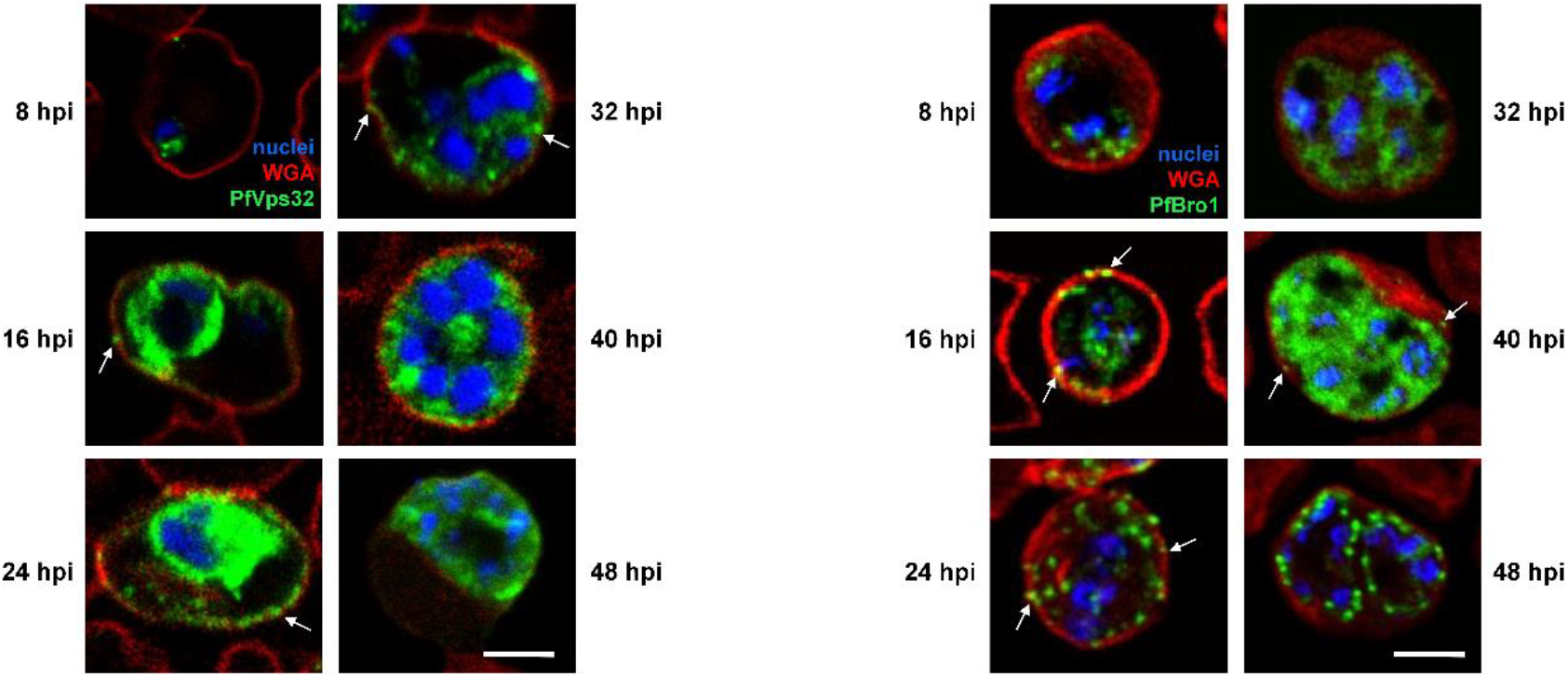
Colocalization of PfBro1-labeled vesicles and WGA in the membrane of pRBCs. Human erythrocytes were infected with *P. falciparum* and fixed at different hpi. PfBro1 (green) and WGA (red) were detected by indirect confocal immunofluorescence microscopy. Cell nuclei were visualized with Hoechst 33342 (blue). Arrows show colocalization events of both proteins in the membrane of the pRBCs. Scale bar: 5 μm

### PfVps32, PfVps60 and PfBro1 are present in extracellular vesicles produced by pRBCs

The results showed above suggested that ESCRT-III proteins could be present in EVs derived from pRBCs. To explore further this hypothesis, we first evaluated by <u>s</u>tochastic <u>o</u>ptical <u>r</u>econstruction <u>m</u>icroscopy (STORM) the presence of PfVps32, PfVps60 and PfBro1 in EVs derived from infected and non-infected RBCs. The sensitivity and resolution of this technique allowed us to detect the protein of interest in single EVs. Parasite proteins were observed in purified EVs from a 3% parasitemia pRBC culture at 40 hpi (Fig 4) and were absent in EVs from non-infected RBCs (data not shown), which confirmed our previous observations and reflected ESCRT-III participation in EVs biogenesis. Anti-GPA antibodies were used to detect EVs from RBC membrane origin. In this case, a significantly higher co-localization of GPA-enriched EVs with PfVps32 was observed when compared with the other two ESCRT proteins studied (Fig 4).

**Fig 4.**
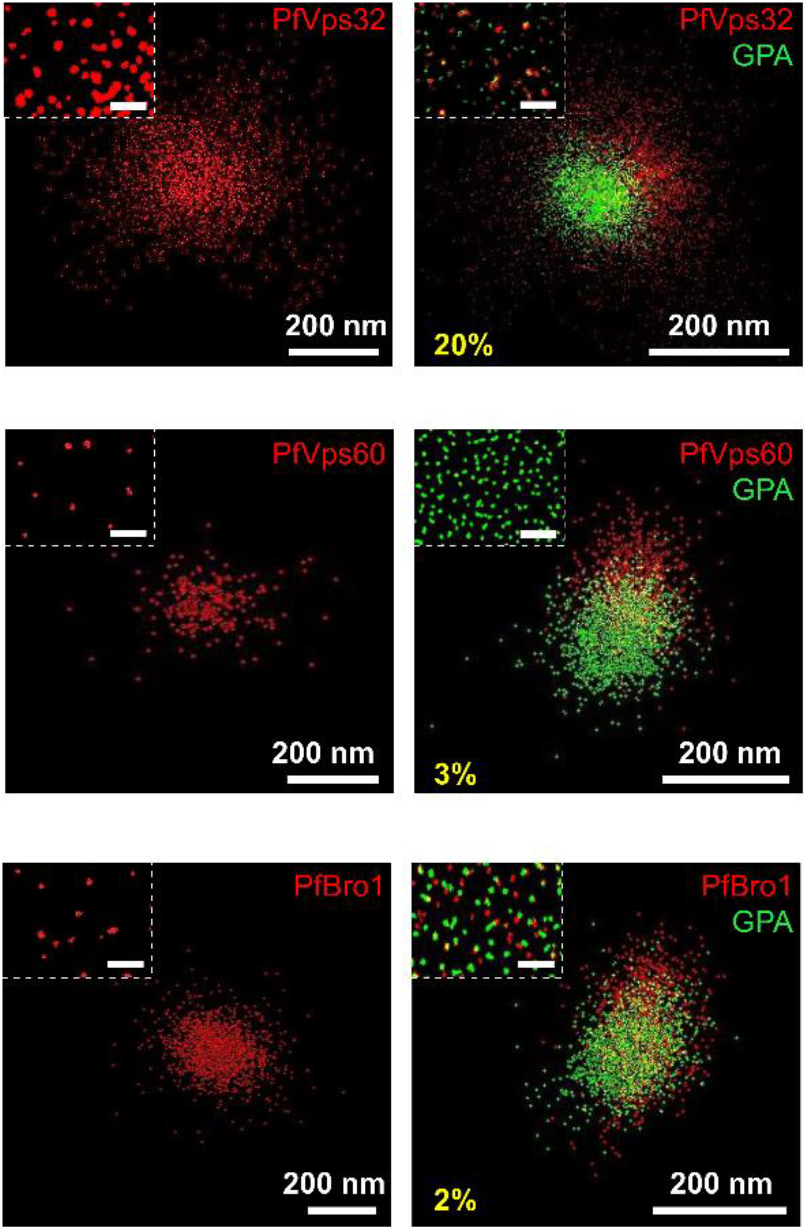
PfVps32, PfVps60 and PfBro1 proteins are present in EVs produced by pRBCs. STORM detection by immunostaining of either PfVps32, PfVps60 or PfBro1 and GPA in purified EVs derived from a 3% parasitemia pRBC culture at 40 hpi. Insets show an enlarged region from the main image. In yellow characters is indicated the percentage of EVs where GPA overlaps with the corresponding parasite protein. Scale bar in insets: 2 μm

As PfBro1 and PfVps60 exhibited a similar protein expression and pattern in the observed EVs, we hypothesize that these vesicles share a common origin. Therefore, the colocalization of individual PfBro1 and PfVps60 molecules in pRBCs was further interrogated by STORM. PfBro1 and PfVps60 were observed to colocalize throughout the whole intraerythrocytic cycle (Fig 5a). PfBro1 was mainly found in large vesicles, whereas PfVps60 exhibited a more homogeneous distribution in the cytoplasm of the parasite and inside the parasitophorous vacuole (PV) (Fig 5a). Interestingly, vesicles labeled with PfBro1 and PfVps60 were detected bound to the surface of non-infected RBCs (Fig 5b, arrows). In all experiments, pre-immune serum used as a control did not display any signal (data not shown).

**Fig 5.**
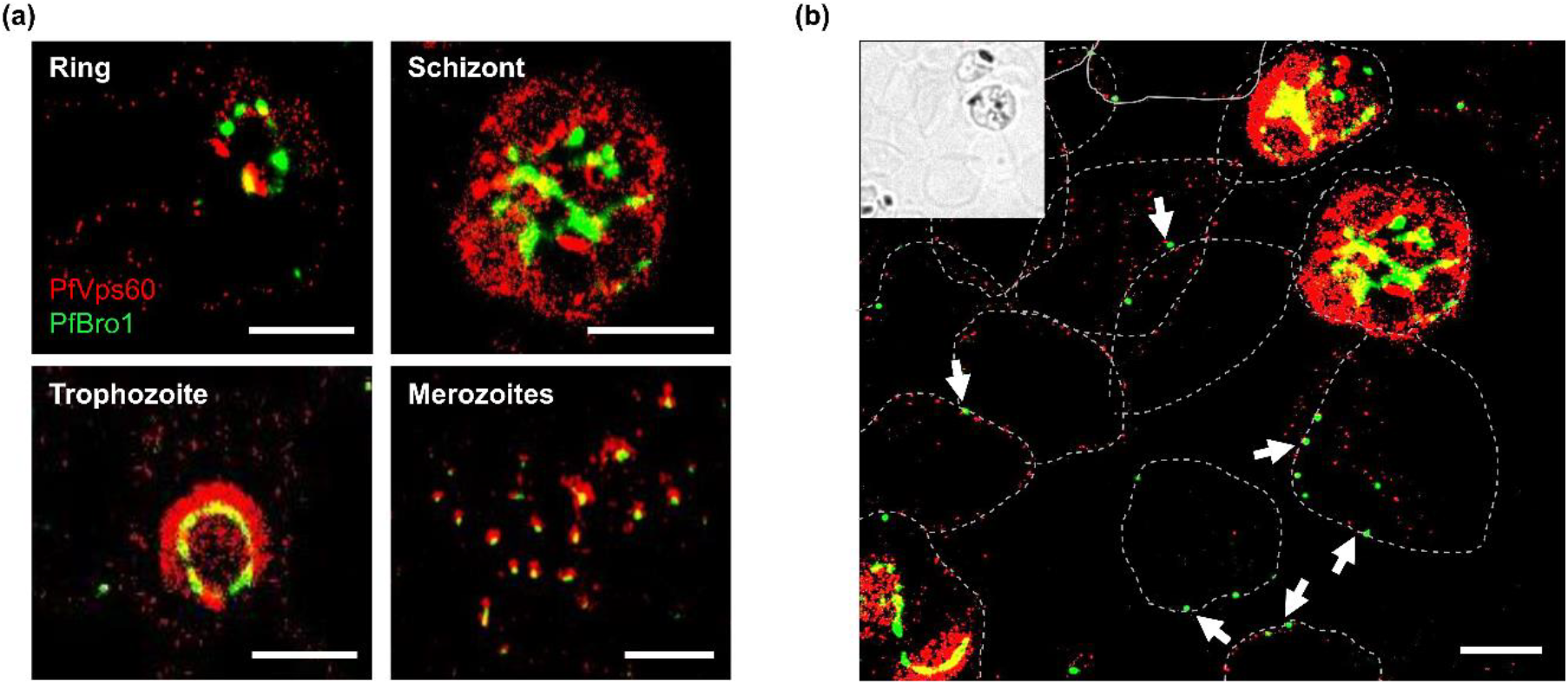
STORM imaging of PfVps60 and PfBro1 colocalization. **(a)** Detection of PfVps60 and PfBro1 in the blood stages of *P. falciparum.* **(b)** Field showing non-infected and infected RBCs. The inset shows a bright field low-resolution image to show the non-infected RBCs. Arrows pinpoint extracellular vesicles bound to non-infected RBCs, whose contours are indicated by dashed lines. Scale bars: 2 μm.

### PfBro1 binds to membranes and recruits both PfVps32 and PfVps60 to trigger bud formation

So far, our results strongly suggested that there is a minimal ESCRT-III machinery participating in the formation of EVs in *P. falciparum*. Due to the fast binding and action of ESCRT-III proteins, it is difficult to assess the function of these proteins in living cells. Other ESCRT-III mechanisms have been studied with the <u>g</u>iant <u>u</u>nilamellar <u>v</u>esicle (GUV) membrane model [33, 34] that allows control of the lipid composition and visualization of the effects of ESCRT-III proteins on membranes by fluorescence microscopy.

To investigate whether *P. falciparum* ESCRT-III-related proteins were able to trigger membrane deformations, GUVs composed by palmitoyl-oleoyl-phosphatidylcholine (POPC) and palmitoyl-oleoyl-phosphatidylserine (POPS) (80:20) were generated to mimic the composition of the inner leaflet from the RBC plasma membrane [35]. We also included the fluorophore 1, 1’-dioctadecyl-3, 3, 3’, 3’-tetramethylindocarbocyanine perchlorate (DiI_C18_) to visualize membrane alterations (see Materials and Methods).

First, we tested the ability of PfBro1 to insert into lipid bilayers using its predicted hydrophobic sequence. When 600 nM of recombinant PfBro1 labeled with Oregon Green^TM^ (OG) 488 (PfBro1-OG488) were incubated with POPC:POPS (80:20) GUVs diluted in an appropriate buffer, the protein inserted into GUVs membranes with a homogenous distribution (Fig 6a). Interestingly, some GUVs with intraluminal buds were occasionally observed at this stage (data not shown). A truncated PfBro1 version lacking its hydrophobic domain (PfBro1t) failed to insert into GUVs membranes (S5 Fig). Incubation in 150 mM NaCl, 25 mM tris-HCl, pH 7.4 (protein buffer) did not affect the GUV morphology (Fig 6a, top panel). After confirming PfBro1 binding to lipid bilayers, we investigated its role as a potential recruiter and activator of ESCRT-III proteins, in particular of Snf7-containg proteins. When POPC:POPS (80:20) GUVs were incubated with 600 nM of unlabeled PfBro1, followed by the addition of 1200 nM of either PfVps32 or PfVps60 labeled with Oregon green 488 (PfVps-OG488) the combination of both proteins induced the formation of intraluminal buds in the GUVs model (Fig 6b-d). The newly formed buds following incubation with PfVps32 were significantly larger (1.95±0.51 μm) than those formed after PfVps60 addition (1.37±0.67 μm) (Fig 6d). Overall, these buds were smaller and more homogeneous in comparison to those where only PfBro1 was used (Fig 6d). Importantly, the buds formed by PfVps60 had a necklace-like arrangement and in the infra-optic range, some tubular structures could be observed (see S1 Video). As the incubation of PfVps32, PfVps60 or PfVps-OG488 alone with GUVs did not produce any detectable membrane changes (Fig 6b, c), we concluded that PfBro1 binds and activates both proteins.

**Fig 6.**
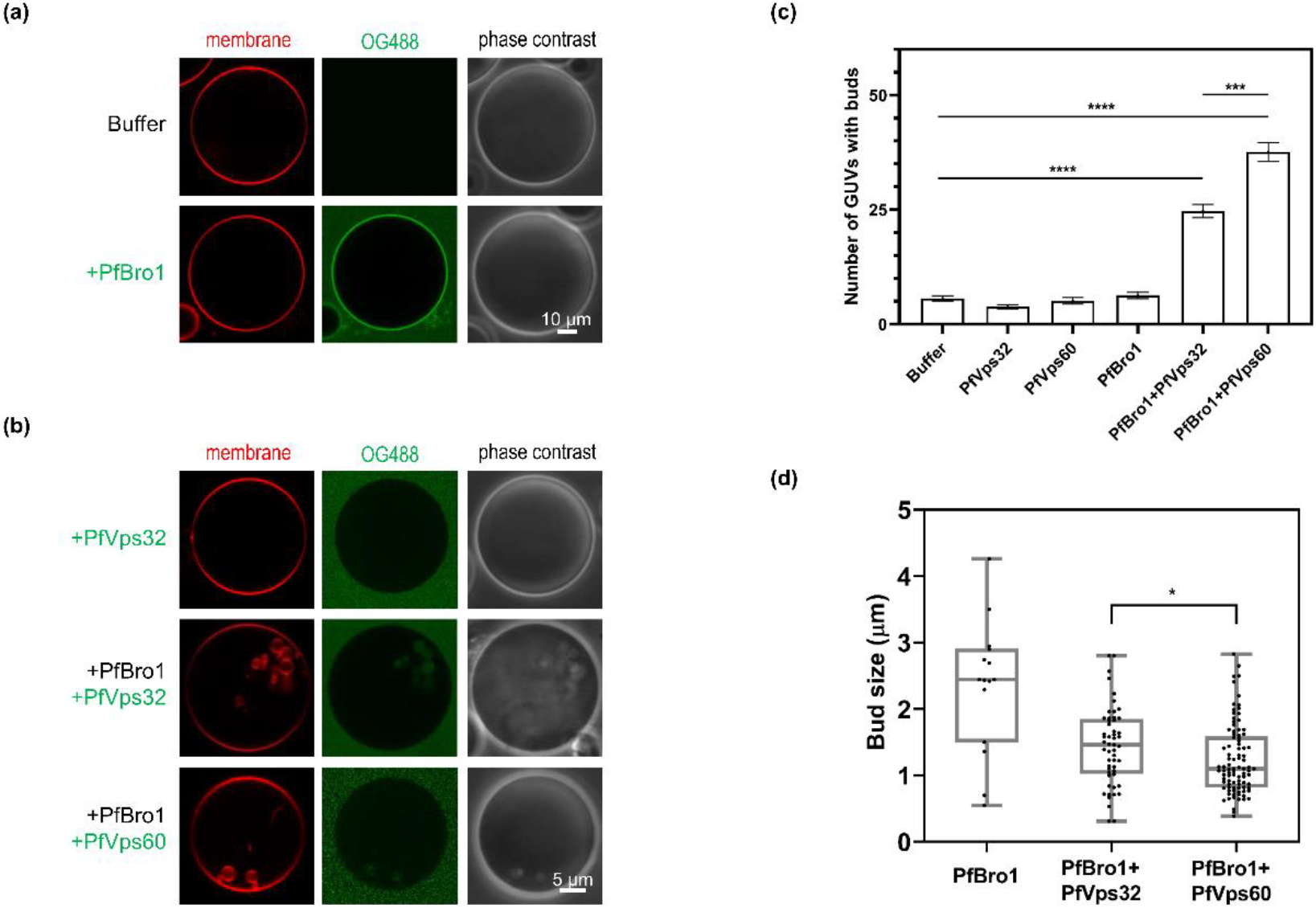
Intraluminal bud formation triggered by ESCRT-III *Plasmodium* proteins. GUVs composed by POPC:POPS (80:20), labeled with DiI_C18_ and diluted 1:2 in protein buffer were incubated with **(a)** 600 nM PfBro1-OG488 with a 1:3 ratio of labeled and unlabeled protein, or **(b)** 600 nM PfBro1 and 1200 nM of either PfVps32 or PfVps60 labeled with OG488 (1:3, labeled: unlabeled) and visualized by fluorescence confocal microscopy. **(c)** Quantification of the number of GUVs with internal buds formed after protein addition. **(d)** Size of buds formed after the addition of the proteins indicated below the graph. Bars represent the mean and standard error of three independent experiments where 50 GUVs were observed. *p* values were determined by Student’s t-test. (*) *p* < 0.05, (***), (**) *p*< 0.01, *p* < 0.001, and (****) *p* < 0.001

### Putative activation of PfVps60 by PfBro1

It is well known that the activation of ESCRT-III subunits occurs after the displacement of the C-terminal domain that is blocking the binding site in the inhibited form of the protein [36]. The rearrangement of this domain has been documented to occur upon binding of activation factors such as Vps20, Vps32 or Bro1 [37]. To check whether this mechanism could be also operating in *P. falciparum*, we performed an *in silico* docking assay using the predicted tertiary structure of the Bro1 domain from PfBro1 and the full-length PfVps60 protein. This pair of proteins was selected because a higher number of GUVs with intraluminal buds was observed for this combination (Fig 6c). Upon binding to the PfBro1 domain, it was predicted that PfVps60 changed from a “closed” to an “open” conformation where the C-terminal domain modified its angle and allowed the exposure of the binding site (S6 Fig). The conformational change predicted by this model is consistent with the GUV protein reconstitution assays presented above.

### PfBro1 and PfVps32 trigger bud formation by direct shedding from the plasma membrane

Although by using the purified recombinant proteins from *P. falciparum* and the GUV model we were able to reconstitute one of the two EV biogenesis pathways described in higher eukaryotic cells (MVB biogenesis; see review in [38]), the mechanism of microvesicles formation by direct shedding from the plasma membrane displays a different topology. To mimic the correct topology involved in this mechanism of EV biogenesis, a microinjection approach was used, where the biotinylated lipid 1,2-distearoyl-sn-glycero-3-phosphoethanolamine-N-[biotinyl(polyethylene glycol)-2000] (DSPE-PEG-biotin) was included in the lipid mixture to form, by a gel assisted method, GUVs containing protein buffer in their lumen. GUVs were harvested and immobilized on an avidin-coated surface to allow their manipulation for injection. It is important to mention that previously to the injection, a z-stack acquisition was performed in the confocal microscope to verify that GUVs lacked alterations in the membrane and that the contact area with the coverslip was not excessively large, which could compromise the assay (see example of selected GUVs in S7 Fig). As protein labeling can compromise protein activity, we used free PEG-fluorescein isothiocyanate (FITC) dye (0.03 mg/ml in protein buffer) to visualize the injection process. The incorporation of this control dye did not produce any detectable alterations in GUVs (see S8 Fig and S2 Video). Upon injection of either PfBro1 or PfVps32 no significant changes were observed in the membrane of the injected GUVs (data not shown), whereas, when a mixture of PfBro1, PfVps32 and PEG-FITC was injected, the formation of extracellular buds was visualized (Fig 7 and S3 Video). Interestingly, the newly formed vesicles remained attached to the mother vesicle moving along its surface (S3 Video). Contrary to the experiments observed in the previous approach (Fig 6), these new buds appeared as single bodies with a homogeneous average size of 0.88 ± .076 μm.

**Fig 7.**
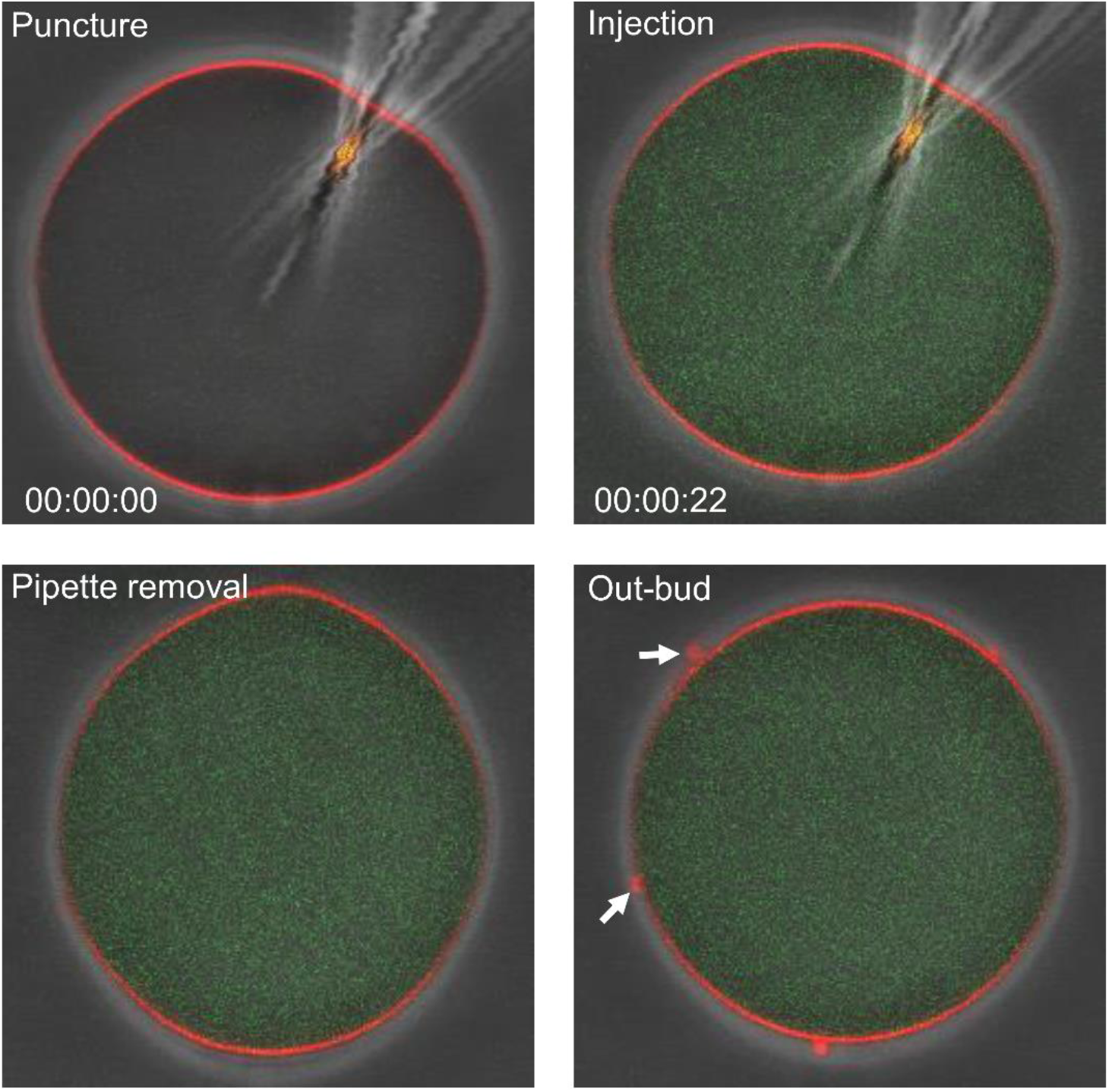
Injection of ESCRT-III *Plasmodium* proteins in GUVs and outward budding. Panels show injection of a mixture of PfBro1 and PfVps32 (1:2) together with PEG-FITC in GUVs composed by POPC:POPS:DSPE-biotin (79:20:1) and labeled with DiI_C18_ (0.1 mol%). Four main events are presented: puncture, injection, pipette removal and generation of outward buds (arrows).

### Disruption of PfVps60 causes a defect in EV production in *P. falciparum*

Next, we evaluated the effect of ESCRT-III machinery inactivation on EV production by *P. falciparum.* While we failed at obtaining a stable strain for the KO of PfVps32 and PfBro1 (probably due to their essential role in the life cycle of the parasite), we succeeded in the establishment of a PfVps60 KO strain by CRISPR/Cas9 gene edition (Fig 8a). Gene silencing and DNA integration were confirmed by diagnostic PCR as shown in Fig 8b. The relative fitness of the generated KO line was evaluated by a growth curve, which showed a slower progression in the KO line compared to its parental line (intraerythrocytic developmental cycle of 52.38 vs. 55.41 h, respectively) (Fig 8c). The suppression of PfVps60 was confirmed by immunofluorescence assays, which indicated the absence of the protein (Fig 8d). In order to study the effects of the PfVps60 KO on the EVs production, we proceed to purify EVs in synchronized cultures after 40 hpi. The number of EVs was significantly reduced in the KO parasites in comparison to the parental line 3D7 (Fig 8e). Accordingly, the total amount of protein exported in the EVs from the KO parasites was also reduced (Fig 8f).

**Fig 8.**
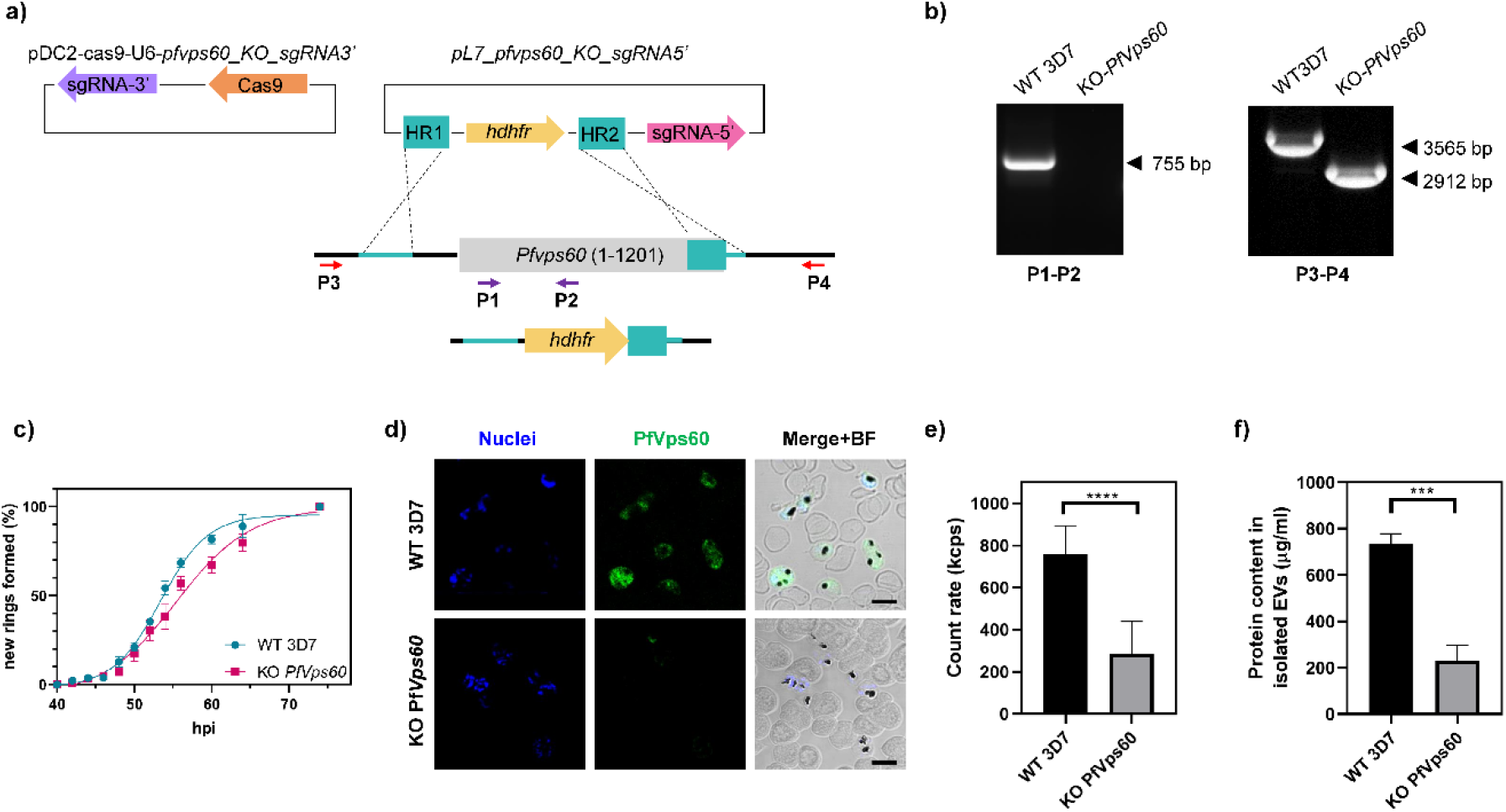
Generation and validation of *PfVps60* KO parasites. **(a)** Schematic of the strategy to generate the transgenic lines using the CRISPR/Cas9 system. Arrows indicate the position of primers used for diagnostic PCR. **(b)** Diagnostic PCR confirmation of the integration of the pL7-*Pfvps60*_KO_sgRNA3’ plasmid at the *Pfvps60* locus. Legend at the bottom indicate the primer pair used for each PCR reaction. Genomic DNA from the WT 3D7 line or the *Pfvps60* KO transgenic line was used. The size of the bands as expected are indicated in the left side. **(c)** Asexual blood cycle duration in the *Pfvps60* KO line compared with its parental 3D7 line. Percentages indicate the proportion of rings observed relative to the total number of rings at the end of the assay. Data was fitted to a sigmoidal curve with variable slope to extract the intraerythrocytic developmental cycle (IDC). **(d)** Human erythrocytes infected with P. falciparum were fixed and PfVps60 was detected by indirect immunofluorescence microscopy. Cell nuclei were visualized with Hoechst 33342 (blue). Fields were merged with bright field (BF) to assess localization. Scale bar: 5 μm. **(e)** Count rate of purified EVs expressed in kilo counts per second (kcps). **(f)** Protein content in purified EVs. Each symbol shows the mean of three different replicates, bars show the SE.

## Discussion

*Plasmodium*-infected erythrocytes increase the release of EVs, which participate in different pathogenic mechanisms (see review in [39]), including cytoadherence during cerebral malaria [40, 41], cell-cell communication between parasites [42, 43], gametocytogenesis induction [42, 43], DNA transport [44] and transfer of drug resistance genes [43]. In higher eukaryotes, the ESCRT-III machinery is involved in both types of EV generation: exosome release and microvesicle budding [45, 46]. However, the mechanisms underlying the release of EVs in *Plasmodium*-infected cells are far from being understood. The *P. falciparum* genome lacks genes encoding for ESCRT-III activating factors, such as Vps20 and ESCRT-II members [28] (S1 Table). Therefore, we hypothesize that there are alternative pathways for ESCRT-III activation in *P. falciparum* and most likely in other intracellular protists such as *Toxoplasma gondii* and *Crytosporidium parvum*, which also lack the aforementioned genes [23]. In *S. cerevisiae* and humans, Bro1 homologues are able to bind directly to the Snf7 domain of Vps32 and CHMP5 (Vps60 homologue) and activate ESCRT-III polymerization on the membranes [24, 30, 47]. In the present study, we show that *P. falciparum* possesses a Bro1 domain-containing protein, PfBro1, capable of activating two SNF7-containing proteins, PfVps32 and PfVps60, with redundant functions. These proteins are expressed throughout the intraerythrocytic cycle. Whereas PfVps60 and PfBro1 exhibited a maximum expression level at 32 hpi, corresponding to the trophozoite stage, PfVps32 had the highest expression in the early stages (8 to 24 hpi). The different expression patterns of these proteins as well as their localization in different subcellular fractions suggest that they display a site- and time-specific action, despite fulfilling similar functions as discussed below.

STORM imaging allowed us to observe the colocalization of PfVps60 and PfBro1 at the asexual blood stages of *P. falciparum* parasites, which suggests their possible association in the parasite. Furthermore, these proteins were detected in EVs at the surface of non-infected erythrocytes, thus suggesting their involvement in a potential pathogenic mechanism that could facilitate invasion of targeted cells. However, further experiments are required to define the role of these EVs in non-infected RBCs.

The identified PfBro1 has a hydrophobic sequence in its C-terminal region that likely allows its insertion into GUVs lipid bilayers, thus making it a good candidate for ESCRT-III recruitment at the membrane. The study of ESCRT-III interactions in living cells is problematic as the association between the different molecular components occurs in a fast manner and the complexes are difficult to obtain. Therefore, *in silico* docking assays were performed and the results showed that PfVps60 can shift from a closed (inactive) to an open (active) conformation upon PfBro1 interaction. Furthermore, we proved that PfBro1 is able to recruit both PfVps32 and PfVps60 to the GUV membrane and activate them, leading to the formation of buds even in the absence of energy, as occurs in the same model with other ESCRT-III homologues [48]. Moreover, the purified proteins were used to recreate the two topologies of EV production (MVB generation and membrane shedding) in GUVs that mimic the composition of the inner leaflet of the erythrocyte’s plasma membrane. As the newly formed buds remained in close contact with the mother vesicle, we hypothesize that more factors are needed to release the nascent vesicle from the membrane. Interestingly, while bud generation is triggered by both PfVps32 and PfVps60 in a similar manner, the nascent vesicles vary in size showing differences depending on the protein added and the employed approach. For instance, PfVps32 lead to significantly bigger buds in comparison to PfVps60 when the proteins are added to the GUV-batch solution. In higher eukaryotes there are several factors governing the size of ESCRT-III-derived buds, including Vps4 disassembly action [49, 50], size of cargo [33] and membrane tension [48]. In the case of *P. falciparum,* whether the size of EVs is regulated by other ESCRT proteins encoded in its genome or by membrane biophysical properties, remains to be explored in the future.

The detection of PfBro1, PfVps32 and PfVps60 in purified EVs from a pRBC culture demonstrates that the observed EVs are of parasite origin and generated by ESCRT-III action. Previous studies found that the isolation method could affect the protein content of *Plasmodium*-derived EVs [42, 43, 51]. In agreement with this observation, our STORM analysis revealed a wide heterogeneity of vesicles, with different apparent size and protein distribution. Importantly, the RBC membrane protein, GPA, was mainly detected in PfVps32-positive EVs, which most probably correspond to EVs derived from the RBC plasma membrane (microvesicles). On the contrary, GPA was much less abundant in PfVps60-positive and PfBro1-enriched EVs, suggesting that their source does not involve the plasma RBC membrane and they are exclusively of parasite origin, probably generated through MVBs formation. These results strongly suggest that both types of EV formation are being carried out in *Plasmodium*-infected RBCs, thus supporting previous observations [42, 43]. Interestingly, GPA has also been detected in *P. falciparum* L-lactate dehydrogenase-carrying EVs which are involved in the control of the parasités proliferation *in vitro* [52]. Whether PfVps32 is also involved in the transport of this protein or others remains to be elucidated. Furthermore, silencing of the *Pfvps60* gene resulted in the reduction of the number of EVs produced during the first 40 hpi, which confirms its participation in EVs biogenesis during *Plasmodium* infection. Although our initial aim was to inactivate the whole ESCRT-III complex, we could not obtain stable KO lines for PfVps32 and PfBro1 proteins, probably due to their involvement in essential processes from the parasite, most likely in cytokinesis as occurs in other eukaryotes [49, 53].

Altogether, our results improve the mechanistic understanding of protein export in *P. falciparum.* Research in this field has mainly focused on the soluble proteins secreted by the parasites, but the mechanism underlying the export of other molecules is still not well understood. Here we describe for the first time a functional ESCRT-III machinery in *P. falciparum* that is involved in the production of EVs in infected erythrocytes, where two redundant proteins, PfVps32 and PfVps60 appear to be time- and site-specific. Due to the low amino acid conservation with their human homologues, as well as their presence throughout the whole intraerythrocytic cycle, the proteins studied here represent a potential target for new therapeutic strategies to control malaria.

## Materials and Methods

### *P. falciparum* culture and synchronization

Unless otherwise indicated, reagents were purchased from Sigma-Aldrich (St. Louis, MO, USA). Asexual stages of *P. falciparum* 3D7 were propagated in group B human erythrocytes at 3% hematocrit using RPMI medium supplemented with 0.5% (w/v) Albumax II (Life Technology, Auckland, New Zealand) and 2 mM L-glutamine. Parasites were maintained at 37 °C under an atmosphere of 5% O_2_, 5% CO_2_ and 90% N_2_. For all experiments, the parasitemia of the culture was maintained between 3 and 5%.

For tight synchronization, the parasite culture was initially synchronized in the ring stage with a 5% sorbitol lysis [54] followed by a second 5% sorbitol lysis after 36 h. Then, 36 h after the second sorbitol, parasites were synchronized in the schizont stages by treatment in 70% Percoll (GE Healthcare, Uppsala, Sweden) density centrifugation at 1,070 × g for 10 min. Finally, after the third synchronization a final 5% sorbitol lysis was done, yielding parasites tightly synchronized at 8 hpi.

*P. falciparum* NF54-*gexp02-tdTomato* gametocytes were induced by choline removal[55] and selected by addition of 50 mM N-acetyl-D-glucosamine[56].

### STORM

A 5% parasitemia RBC culture was prepared for super-resolution microscopy as described in[57]. Briefly, a μ-Slide 8 well chamber slide (ibidi) was coated for 20 min at 37°C with 50 mg/ml concanavalin A. Then, wells were rinsed with pre-warmed phosphate buffered saline (PBS) before parasite seeding. Infected RBCs were washed twice with PBS and deposited into the wells. Cells were incubated for 10 min at 37 °C and unbound RBCs were washed away with three PBS rinses. Seeded RBCs were fixed with pre-warmed 4% paraformaldehyde at 37 °C for 20 min. After this time, cells were washed with PBS and then, incubated with antibodies α-PfVps32-Alexa Fluor 488, α-PfVps60-Alexa Fluor 488 (1:500) and α-PfBro1-Alexa Fluor 647 (1:1000). Finally, nuclei were counterstained with Hoechst 33342 (2 μg/ml).

Before STORM acquisition, the buffer was exchanged to OxEA buffer (3% V/V oxyrase, 100 μM DL-lactate, 100 mM β-mercaptoethylamine, dissolved in 1× PBS, pH 8.4) [58]. STORM images were acquired using a Nikon N-STORM system configured for total internal reflection fluorescence imaging. Excitation inclination was tuned to adjust focus and to maximize the signal-to-noise ratio. Alexa Fluor 647 and 488 were excited, respectively, illuminating the sample with 647 nm and 488 nm laser lines built into the microscope. Fluorescence was collected by means of a Nikon 100×, 1.4 NA oil immersion objective and passed through a quad-band-pass dichroic filter (97335 Nikon). 20,000 frames at 50 Hz were acquired for each channel. Images were recorded onto a 256×256 pixel region (pixel size 160 nm) of a CMOS camera. STORM images were analyzed with the STORM module of the NIS element Nikon software.

### Reconstitution of ESCRT-III in GUVs

GUVs containing POPC, POPS, and the fluorophore DiI_C18_ (Invitrogen, CA, USA) (80:20:0.1) were prepared in 600 mM sucrose as described previously [34]. Briefly, the lipid mix was spread on tin oxide-coated glass slides, and electro-swelling was performed for 1 h at room temperature (RT) at 1.2 V, and 10 Hz. All lipids were obtained from Avanti Polar Lipids (Alabaster, IL, USA).

For PfBro1 binding assays, GUVs were harvested and diluted 1:1 with 2 folds’ protein buffer (50 mM tris-HCl, 300 mM NaCl, pH 7.4). After 10 min of equilibration, GUVs were incubated with 600 nM of either OG488-PfBro1 or OG488-PfBro1t. For PfVps32 recruitment, equilibrated GUVs were incubated with 600 nM of PfBro1 and 1200 nM of either PfVps32 or PfVps60 with at least 10 min of incubation at RT between the additions of each protein. Images were acquired with a Leica TCS SP5 confocal microscope (Mannheim, Germany). DiI_C18_ was excited with a 561 nm laser and OG488 with a 488 nm line of an Argon laser. To avoid crosstalk between the different fluorescence signals, a sequential scanning was performed. All experiments shown in the same figure were done with the same GUV batch for comparability. Each experiment was repeated on at least three separate occasions with different batches of GUVs.

### Femtoliter injection

A lipid mixture of POPC, POPS, DSPE-PEG-biotin, and DPPE-rhodamine (78.9:20:1:0.1 mol%) was prepared in chloroform. GUVs filled with protein buffer were grown by the gel-assisted method. Briefly, a 5% (w/w) polyvinyl alcohol (PVA) solution was prepared in protein buffer (25 mM tris-HCl, pH 7.4, 150 mM NaCl). The PVA solution was spread on a microscope coverslip and then dried for at least 30 min at 50 °C. 10-15 μl of lipids dissolved in chloroform (1 mg/ml) were spread on the dried PVA film and placed under vacuum for 1 h to eliminate the solvent. A chamber was formed with a homemade Teflon® spacer sandwiched between two glasses and filled with protein buffer for 10 min at RT. Then, GUVs were harvested by gentle tapping on the bottom of the chamber and collected using a micropipette without touching the PVA film to avoid sample contamination. To immobilize GUVs, cleaned coverslips were incubated for 20 min at RT with a 1:1 mixture of 1 mg/ml BSA-biotin, 1 mg/ml BSA (both diluted in protein buffer to maintain osmolarity). After incubation, coverslips were washed with distilled water and incubated with 0.005 mg/ml avidin. Subsequently, slides were washed and dried with N_2_. These coverslips were used to assemble a homemade observation chamber using a Teflon® spacer, GUVs were deposited and let to settle down for at least 10 min.

The micropipettes used to perform the injection were fabricated from thin wall borosilicate capillaries glass with filament (Harvard Apparatus, Holliston, MA, USA) in a pipette puller (Sutter Instruments, Novato, CA, USA) to obtain bee-needle type tips. For the injection experiments, immobilized GUVs were imaged under a Leica TCS SP5 confocal microscope. The micropipette was placed on a mechanical holder attached to a micromanipulator (Sutter Instruments) and then connected to a Femtojet microinjector set (Eppendorf). Injection was performed in a 15° angle, using a pressure of injection of 150 hPa, time of injection of 5.0 s and a compensation pressure of 1 hPa. The solution injected corresponded to a 4× protein mixture stock (2.4 nM PfBro1 and 4.8 nM of either PfVps32 or PfVps60 dissolved in 1× buffer) and 0.03 mg/ml PEG-FITC to monitor injection.

### Generation of *Pfvps60* KO strain

Homology regions (HR) of the 5’UTR (HR1, spanning positions −762 to −243 from the *Pfvps60* start codon) and 3’UTR (HR2, spanning positions 1,012 to 1,553 from the *Pfvps60* start codon) were PCR amplified using genomic DNA purified from a *P. falciparum* 3D7 culture synchronized at late stages. Primers used for PCR amplification are listed in table S2. The generated HR1 and HR2 were cloned by ligation using restriction sites *SpeI* and *AflII* (HR1), and *EcoRI* and *NcoI* (HR2) into a modified pL6-*egfp* donor plasmid [59] in which the *yfcu* cassette had been removed [60]. The single guide RNA (sgRNA) specific for the *Pfvps60* gene and targeting the sequence near the 5’ end (sgRNA 5’, position −225, −206) was generated by cloning annealed oligonucleotides into the *BtgZI* site to generate the pL7-*pfvps60*_KO_sgRNA5’ plasmid. On the other hand, the pDC2-Cas9-U6-h*dhfr* vector [61] was modified by cloning a sgRNA specific for the sequence near the 3’ end (sgRNA 3’, position 980, 999) into the BtgZI site of this plasmid to generate the pDC2-Cas9-U6*-pfvps60*_KO_sgRNA3’ plasmid. All guides were cloned using the In-Fusion system (Clontech, Japan).

For transfection of 3D7 rings, 60 μg of circular pDC2-Cas9-U6-h*dhfr-* pfvps60_KO_sgRNA3’ plasmid and 30 μg of linearized (with *PvuI*) donor plasmid were precipitated, washed and resuspended in 30 μl of sterile 10 mM Tris, 1 mM EDTA (TE) buffer. Then, plasmids were diluted in 370 μl of Cytomix buffer (120 mM KCl, 0.15 mM CaCl_2_, 10mM K_2_HPO_4_/KH_2_PO_4_, 25 mM Hepes, 2 mM EGTA, 5 mM MgCl_2_, pH 7.6) and introduced into parasites by electroporation using a Bio-Rad Gene Pulser Xcell^TM^ system, at 310 V, 950 μF of capacitance and without resistance. Electroporated parasites were carefully recovered and resuspended in RMPI medium supplemented with 0.5% (w/v) Albumax II and 2 mM L-glutamine. Twenty-four hours after transfection, cultures were selected with 10 nM WR99210 for 4 consecutive days [62]. To validate the integration of the plasmids, a diagnostic PCR analysis was performed using LA Taq® DNA polymerase (Takara, Japan), the primers listed in table S2 and gDNA obtained from the *Pfvps60* KO strain and compared with the WT 3D7 strain. The fitness of the generated line compared with the parental 3D7 line, was evaluated by calculating the percentage of newly formed rings in tightly synchronized cultures. Initial parasitemia was determined at ~18 hpi, then rings parasitemia was determined at different time points within the period were most schizont bursting and reinvasion events occurred (44 to 62 hpi). The final point was 74 hpi when all viable schizonts had burst [60]. Data points were determined by the proportion of rings relative to the total number of rings at the end of the assay. Data was fitted to a sigmoidal dose-response curve and the time to generate 50% or the rings in each population was determined [63].

## Acknowledgments

We thank Harvie Portugaliza and Alfred Cortés for the *P. falciparum* NF54 gexp02-tdTomato transgenic line. We are also grateful to Dr. Elisabet Tintó-Font for donating the plasmids that were used for our CRISPR/Cas9 constructs. Authors acknowledge Dr. Igor Florez-Sarasa for critical reading of this manuscript.

