## Supplementary Information for "The ESCRT-III machinery participates in the production of extracellular vesicles and protein export during *Plasmodium falciparum* infection"

### Supporting information

| *Plasmodium falciparum* | | *Homo sapiens* | | | | | *Saccharomyces cerevisiae* | | | | | |
|---|---|---|---|---|---|---|---|---|---|---|---|---|
| **Predicted protein** | **Accession number** | **Protein** | **Accession number** | **E- value** | **S (%)** | **I (%)** | **Protein** | **Accession number** | **E-value** | **S (%)** | **I (%)** | **Function** |
| PfVps32 | Q8IL52 | CHMP4A | Q9BY43 | - | - | - | Snf7 | P39929 | 4.00E-06 | 60 | 19 | Binds to Vps20/Bro1, forms concentric polymers that help to close the neck of the nascent vesicle. |
| | | CHMP4B | Q9H444 | - | - | - | | | | | | |
| | | CHMP4C | Q96CF2 | - | - | - | | | | | | |
| PfVps60 | A0A144A0N0 | CHMP5 | Q9NZZ3 | 4.00 E-11 | 79 | 23 | Vps60 | Q03390 | 6.00E-04 | 78 | 21 | Binds to Brox and Vta1, mediates Vps4 activity. |
| PfVps2 | Q8IAZ9 | CHMP2a | O43633 | 2.00E-24 | 65 | 32 | Vps2 | P36108 | 1.00 E-24 | 63 | 33 | Forms filaments together with Vps24 and promote |
| | | CHMP2b | Q9UQN3 | 9.00E-15 | 64 | 26 | | | | | | |
| PfVps46 | Q8I388 | CHMP1a | Q9HD42 | 7.00E-24 | 85 | 29 | Vps46 | P69771 | 7.00E-17 | 58 | 30 | Controls membrane association and activity of Vps4 |
| | | CHMP1b | Q7LBR1 | 3.00E-18 | 56 | 31 | | | | | | |

**S1 Table. SNF-7 domain-containing proteins encoded in the *P. falciparum* genome**. Modified from[23]. Percentages of similarity (S), identity (I) and expectation value (E-value) relative to P. falciparum proteins were determined using the Expert Protein Analysis Systems (ExPASy) Proteomics Server by the NCBI BLAST service program.

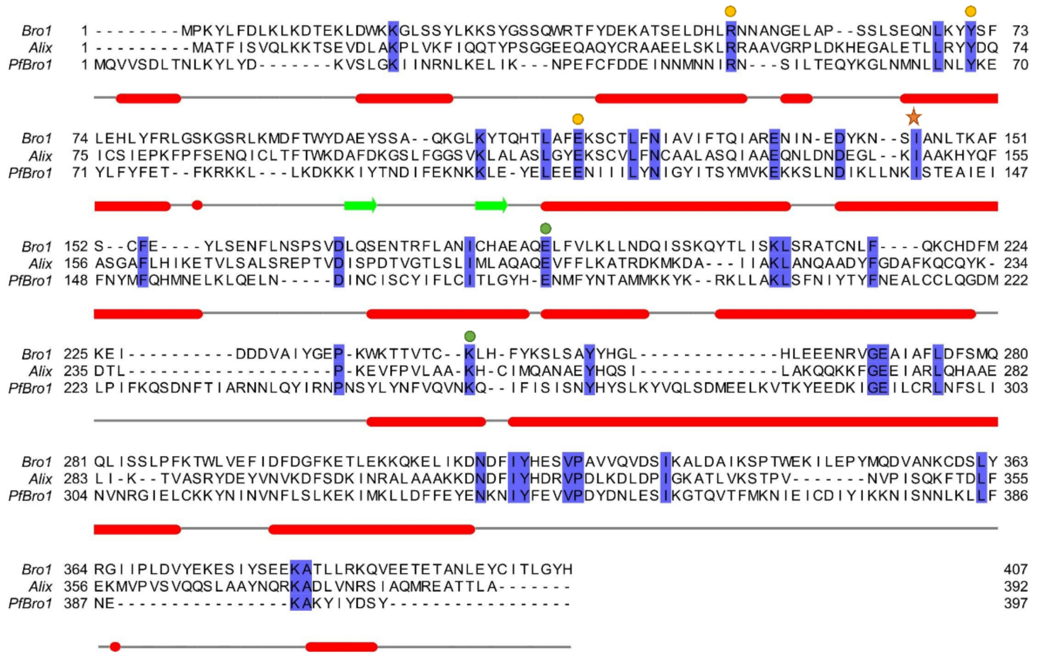

**S1 Fig. Multiple sequence alignment for Bro1-homologues in yeast (Bro1), *P. falciparum* (PfBro1) and humans (ALIX).** Conserved residues are shadowed in violet. Conserved amino acids present in Bro1-containing proteins are indicated with colored circles, yellow for the polar cluster I and green for the polar cluster II. A key isoleucine involved in Vps32 binding is indicated with a star. The secondary structure for PfBro1 is displayed below the sequences, alpha helices represented in red and beta-sheets in green. Sequence alignments were performed with Clustal Omega and edited in Jalview.

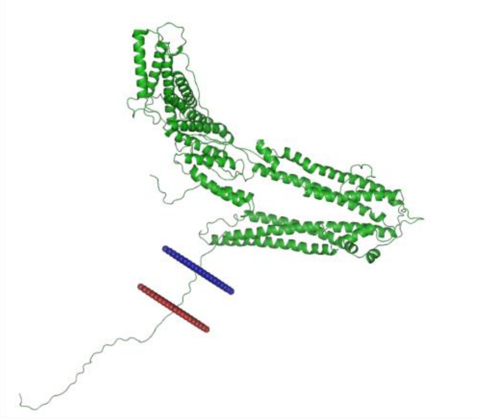

**S2 Fig. Tertiary prediction and membrane orientation of PfBro1.** Outer membrane is colored in red, inner membrane in blue. The structure was generated using the Phyre2 server and the OPM database.

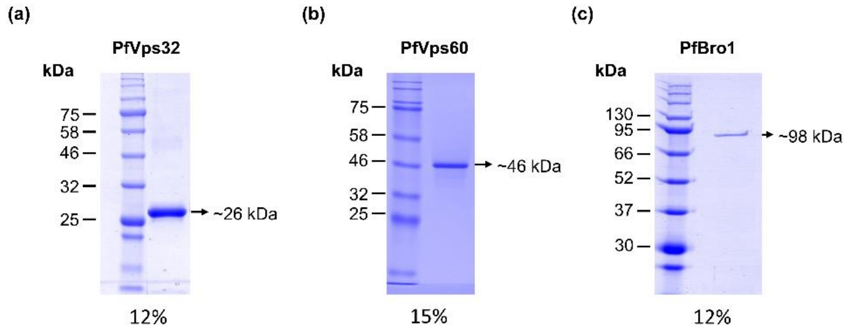

**S3 Fig. Purification of proteins.** 12% and 15% SDS-PAGE stained with Coomassie Blue showing the purified fractions of **(a)** PfVps32, **(b)** PfVps60 and **(c)** PfBro1 that were used for this study.

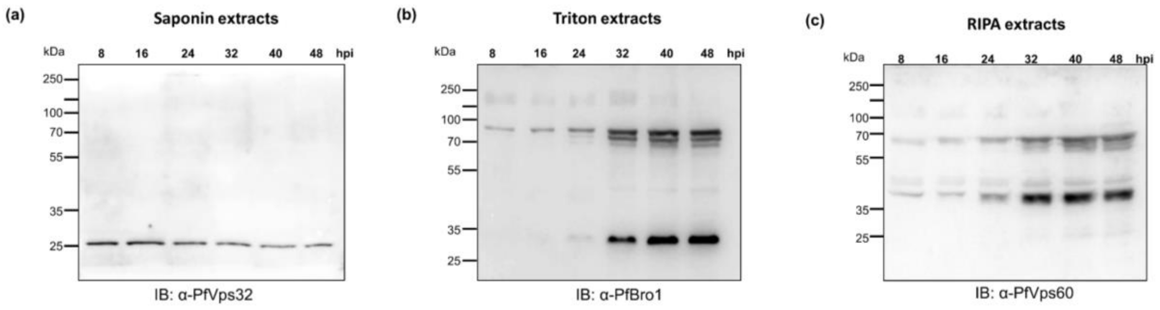


**S4 Fig. Expression of PfVps32, PfVps60 and PfBro1 during the *P. falciparum* intraerythrocytic cycle.** Uncropped Western blot analysis of *P. falciparum* in **(a)** saponin, **(b)** Triton X-100 or **(c)** RIPA buffer protein extracts at different hpi presented in Fig 1.

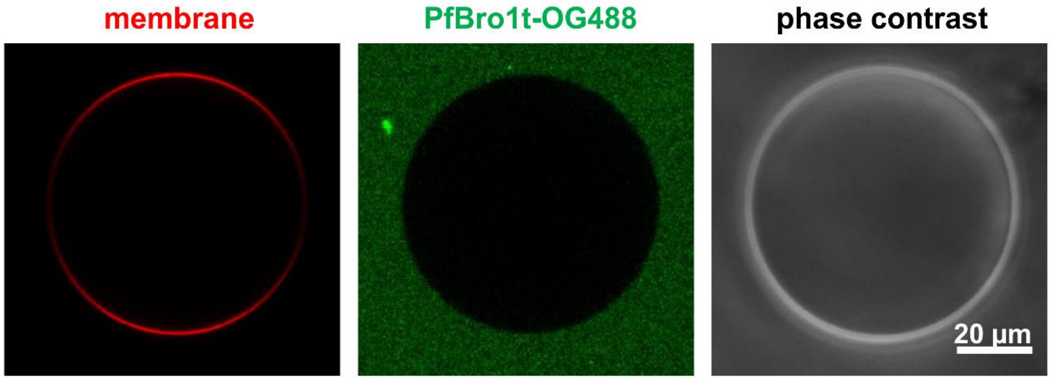


**S5 Fig. Incubation of PfBro1t-OG488 with GUVs.** POPC:POPS (80:20) GUVs labeled with $DiI_{C18}$ and diluted in protein buffer were incubated with 600 nM of PfBro1t in a 1:3 ratio (labeled:unlabeled protein).

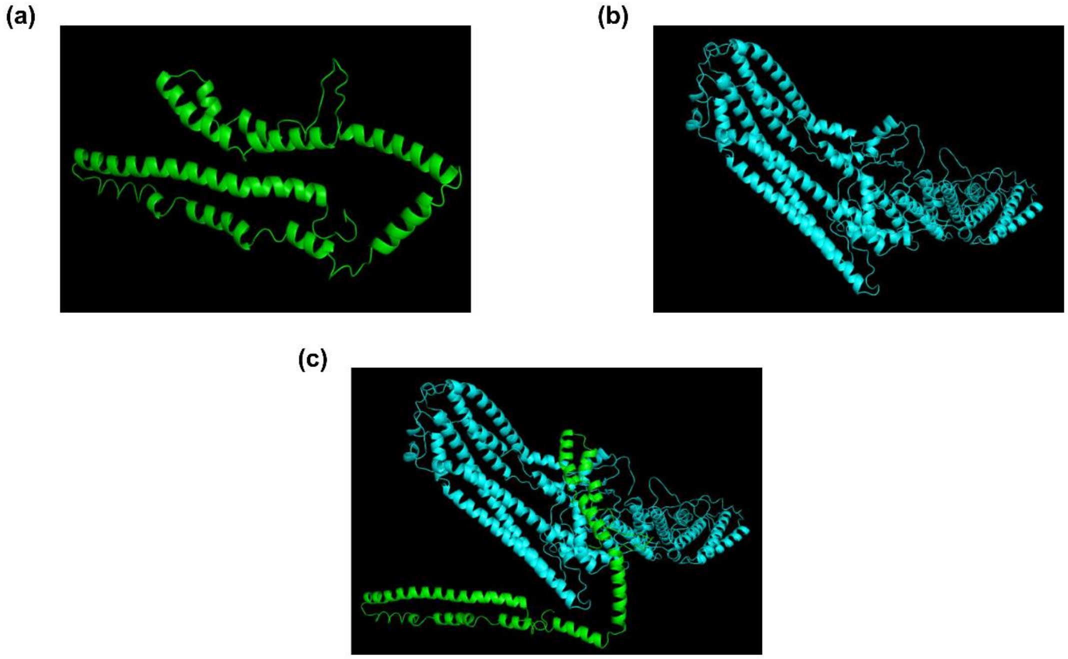


**S6 Fig. Protein docking of PfBro1 and PfVps60.** Predicted structure of **(a)** PfVps60 in its auto-inhibited form and **(b)** PfBro1 domain. **(c)** Protein docking simulation showing the PfVps60 “opening”. All images were generated using PyMOL.

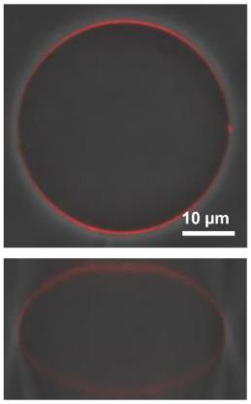

**S7 Fig. Representative images of a GUV selected for injection.** Panels show the top and side view of a typical vesicle selected to perform protein injection.

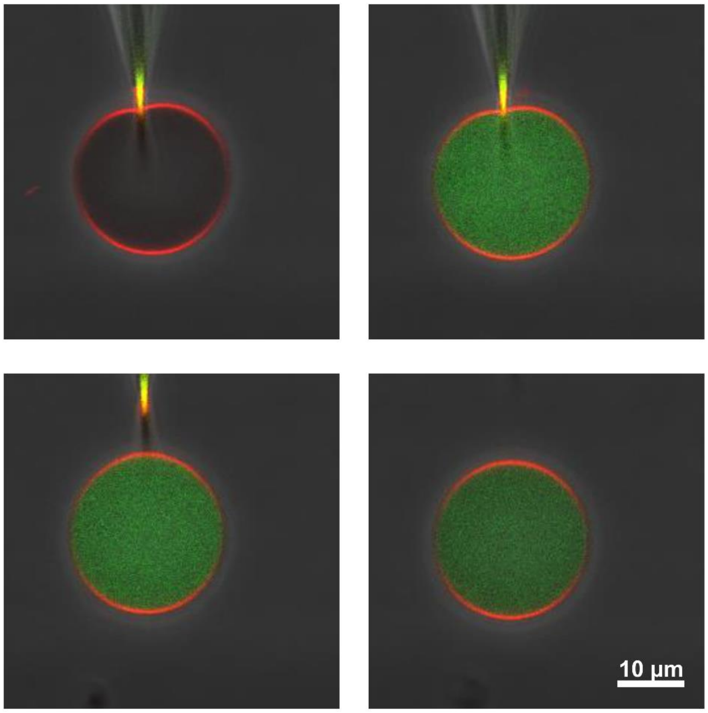


**S8 Fig. Femtoliter injection of PEG-FITC in protein buffer.** GUVs composed by POPC:POPS:DSPE-biotin (79:20:1) and labeled with DPPE-rhodamine (0.1 mol%) were grown in a PVA-substrate using protein buffer, harvested after 10 min and deposited on an avidin-coated coverslip, and injected with PEG-FITC. No alterations were observed up to 5 min after injection.

**S2 table. Primers used for CRISPR/Cas9 gene edition.**

**S1 Video.** POPC:POPS (80:20) GUVs labeled with $DiI_{C18}$ (0.1 mol%) and incubated with 600 nM PfBro1 and 1200 nM PfVps60. Intraluminal and interconnected buds of different sizes are observed.

**S2 Video.** Injection of POPC:POPS:DSPE-biotin (79:20:1) GUVs labeled with DPPE-rhodamine (0.1 mol%) with free PEG-FITC. The video is accelerated 4.5×.

**S3 Video.** Injection of POPC:POPS:DSPE-biotin (79:20:1) GUVs labeled with DPPE-rhodamine (0.1 mol%) with a mixture of PfBro1:PfVps32 (1:2) together with PEG-FITC. The video is accelerated 4.5×.

#### Supplemental Materials and Methods

Unless otherwise indicated, reagents were purchased from Sigma-Aldrich (St. Louis, MO, USA).

##### *In silico* analysis of PfBro1

The Bro1 protein domain was searched in the *P. falciparum* genome database (http//www.plasmodb.org). Comparison between putative PfBro1 and human and yeast orthologues was determined using the Expert Protein Analysis Systems (ExPASy) Proteomics Server by the NCBI BLAST service program. Sequence alignments were generated using the Clustal Omega program[1] and edited in Jalview[2].

##### Cloning and expression of PfVps32, PfBro1 and PfBro1t

The proteins PfVps32A, PfVps43B and PfBro1 (Uniprot accession numbers A0A144A0N0, Q8IL52, and Q8I5H5, respectively) were expressed from a codon-optimized gene (Genscript, Leiden, The Netherlands) inserted into pGEX-6P-1 (GE-Healthcare), as a fusion protein with a glutathione-S-transferase (GST) tag linked to the N terminus of the proteins via a preScission protease site. To generate truncated *PfBro1t*, the first 397 residues from *PfBro1* were PCR-amplified using plasmid cDNA as template and specific primers (CCGGATCCGATGCAAGTGGTTAGCGACCTGACCAATCTGAAATAC and CCGTCGACGTAGCTATCATAGATGTATTTCGCCTTCTCG), which introduced unique *Bam*HI and *Sal*I (underlined) sites in the sense and antisense primers, respectively. The *PfBro1t* gene was cloned into the pJET1.2/blunt plasmid (ThermoFisher), accordingly to manufacturer's instructions. Then, *PfBro1t* was subcloned into the pGEX-6P-1 plasmid.

The proteins GST-PfVps32A, GST-PfVps32B GST-PfBro1 and GST-PfBro1t were expressed in *Escherichia coli* C43(DE3) for 16 h at 20 °C in Terrific Broth (TB) medium. Cells were lysed by sonication in lysis buffer (50 mM tris-HCl, pH 8.0, 300 mM NaCl, 1 mM EDTA, 1 mM PEFABLOC SC (Roche, Mannheim, Germany), and 2 mg/ml lysozyme). Cleared lysates were applied to glutathione Sepharose™ 4B resin (GE healthcare) for at least 5 h at 4 °C under gentle stirring. After this time, the resin was washed 4× with PBS and incubated with PreScission protease in cleavage buffer (50 mM tris-HCl, pH 7.5, 150 mM NaCl, 1 mM EDTA, 1 mM dithiothreitol) for 4 h at 4 °C. Following incubation, the GST-free proteins were recovered after centrifugation of the resin. The eluted proteins were concentrated and applied to a Superdex™ 200 10/300 GL (GE Healthcare) and separated in PBS, pH 7.4. The proteins were labeled overnight at 4 °C with Oregon Green™ 488 succinimidyl ester (Invitrogen). Next, free fluorophore and unlabeled fractions were removed on a Superdex 200 10/300 gel filtration column in 50 mM tris-HCl, pH 7.4, 300 mM NaCl. The pooled Superdex 200 fractions were confirmed by sodium dodecyl sulfate-polyacrylamide gel electrophoresis (SDS-PAGE) stained with Coomassie Blue, concentrated and snap frozen in liquid nitrogen in small aliquots until use.

**Labeling of recombinant proteins**

The purified recombinant proteins were labelled using Oregon Green (OG) 488 (Molecular Probes-Thermo Fisher, Eugene, OR, USA) following the manufacturer's protocol. The labelled and unlabelled proteins were separated by size exclusion chromatography with a Superdex 200 16/600 column (GE-healthcare, Freiburg, Germany) connected to an Äkta-Purifier FPLC (GE-healthcare, Freiburg, Germany). The degree of labelling was assessed according to manufacturer's instructions. In all cases, a 1:3 ratio of labelled:unlabelled proteins was used to maintain activity.

**Generation and labeling of polyclonal antibodies**

PfBro1 (400 µg) or PfVps32A or B (413 µg) were emulsified in complete Freund's adjuvant (1:1) and inoculated into *New Zealand* male rabbits. Three more doses of 200 µg of each protein emulsified in incomplete Freund's adjuvant were injected at 21-day intervals and animals were bled to obtain α-PfVps32 and α-PfBro1 polyclonal antibodies. Polyclonal IgGs were purified using protein A sepharose affinity chromatography. Pre-immune serum was obtained before immunization in all cases.

The labeling of the antibodies for the co-localization experiments was performed using either Alexa Fluor™ 647 NHS ester, Alexa Fluor™ 488 NHS ester or NHS-fluorescein (ThermoScientific) accordingly to manufacturer's instructions. Briefly, 0.1 ml of 1 M $NaHCO_3$ were added to 5 mg of antibody solution, and the mixture was incubated with the different fluorophores overnight at 4 °C. The free dye was removed using a desalting PD-10 column and PBS.

**Subcellular protein extraction**

For the analysis of PfBro1 and PfVps32 homologues throughout the whole intraerythrocytic cycle of *P. falciparum*, tightly synchronized cultures (8, 16, 24, 32, 40 and 48 hpi) were obtained as described before. Then, samples from each time point underwent differential detergent fractionation as previously described[3]. Briefly, infected RBC pellets were washed with PBS-complete buffer (PBS supplemented with 1× cOmplete™ protease inhibitor cocktail, Roche). Then, pellets were suspended in 6× volumes of 0.15% saponin for 10 min at 4 °C and centrifuged (10,000× g, 15 min, 4 °C); the resulting supernatant contained cytosolic soluble proteins. The remaining pellet was washed with PBS-complete and incubated in 1% Triton X-100 for 30 min at 4 °C. Again, samples were centrifuged (20,000× g, 30 min, 4 °C) and the new supernatant containing membrane and organelle proteins was recovered. Finally, Triton-insoluble pellets were incubated for 30 min with 100 µl of RIPA buffer (40 mM tris-HCl, pH 7.4, 150 mM NaCl, 2 mM EDTA, 10% glycerol, 1% Triton X-100, 0.5% sodium deoxycholate, 0.2% SDS and 1× cOmplete™ protease inhibitor cocktail) under continuous and vigorous shaking at 4 °C. Extracts were sonicated for 2 s at 40% amplitude and incubated on ice for 15 min. Finally, extracts were centrifuged at 20,000× g to eliminate undissolved molecular debris. The resulting supernatant contained nuclear and detergent-resistant cytoskeletal/matrix proteins.

**Electrophoresis and Western blot**

Protein extracts (20 µg) of each time point were separated in a 12% SDS-PAGE, transferred to a polyvinylidene difluoride membrane (BioRad, Hercules, CA, USA) and blocked with 5% non-fat milk dissolved in tris buffered saline (TBS: 50 mM tris-HCL, pH 7.6, 150 mM NaCl) containing 0.5% Tween 20. Membranes were probed with either rabbit α-PfVps32A (1:10,000), rabbit α-PfVps32B (1:12,000), rabbit α-PfBro1 (1:8,000), rabbit α-PfHSP70 (StressMarq, 1:10,000) or mouse α-spectrin alpha/beta (1:10,000) antibodies for 3 h at RT. After this time, membranes were washed and incubated for 1 h at RT with α-rabbit or α-mouse HRP-labeled secondary antibodies (Abcam, 1:10,000) and developed with ECL Prime Western blotting detection reagent (GE Healthcare). Western blot analysis was performed using a LAS-4000 image reader (GE Healthcare). Densitometry analysis was done in Fiji software[4]. For the protein densitometry data, statistical analysis was performed with SPSS for Windows 25.0. The Tukey post hoc test was used to determine statistically significant differences.

**Light microscopy and image processing**

For indirect immunofluorescence detection of PfVps32 homologues, PfBro1, ferredoxin-NADP reductase (α-apicoplast) and WGA, thin smears of synchronized cultures at different hpi were air-dried and fixed with a 9:1 mixture of acetone:methanol for 2 min at RT. Slides were thoroughly washed with PBS and incubated for 1 h at RT with the primary antibodies α-PfBro1, α-PfVps32 and α-ferredoxin-NADP reductase (Abcam) diluted 1:200 in PBS supplemented with 0.75% BSA. After washing with PBS, Alexa Fluor 488- or Alexa Fluor 647-labeled secondary α-rabbit IgG antibodies were used at 1:500 dilution. For colocalization assays, either α-PfBro1 or α-PfVps32 previously labeled with Alexa Fluor 647 were used at 1:500 dilutions. Lectins present on the RBC surface were stained using WGA-rhodamine (Invitrogen, 1:500). Nuclei were counterstained with Hoechst 33342 (1:5,000). All preparations were preserved using ProLong$^{TM}$ gold antifade reagent (Invitrogen) and images were collected with a Leica TCS SP5 confocal microscope (Mannheim, Germany) using a 100× oil immersion objective. All immunofluorescence assays were repeated three times. To quantify colocalization, images were analyzed using the Just Another Colocalization Plugin (JACoP)[5] in the Fiji software[4].

#### EVs purification

For purification of EVs, medium was collected from RBC suspensions of 3% parasitemia pRBC cultures at 40 hpi. EV isolation was performed adapting a previously described protocol[6] by centrifugation and ultrafiltration followed by size exclusion chromatography using Sepharose CL-4B. Briefly, samples were prepared by sequential centrifugations of conditioned medium to remove large aggregates. Therefore, cultures were centrifuged for 10 min at 400× g, the cell pellet was discarded, and supernatant was further centrifuged at 2,000× g for 10 min twice. In both cases, a small pellet was discarded. Finally, the supernatant was placed in an Amicon$^{TM}$ Ultra-15 centrifugal filter (100 kDa cut-off, Millipore-Merck, Cork, Ireland) and centrifuged for 20 min at 3,400× g. One ml of the resulting concentrated solution was collected and transferred to a 10 ml homemade Sepharose CL-4B column previously equilibrated with PBS. Finally, EV purification was performed by gravity flow at RT and 0.5-ml fractions were collected. EVs were enriched in fractions 8 and 9.

#### EVs imaging

For STORM visualization of purified EVs, the protocol for super-resolution microscopy of vaccinia virus particles described by Gray & Albrecht[7] was followed. Briefly, a clean coverslip was washed 3 times with ethanol, acetone and deionized water sequentially. Then, coverslips were sonicated in 1 M KOH for 20 min at RT, washed thoroughly with water and placed in a 12-well plate. Next, the purified EVs were sonicated for 30 s to avoid aggregation, diluted in 1 mM tris-HCl, pH 9.0, and deposited on the cleaned coverslips. EVs were left to adhere for 60 min at RT and the rest of the solution was carefully removed with a pipette. The bound EVs were then fixed with 4% paraformaldehyde for 15 min at RT, washed and incubated in quenching buffer (0.25% $NH_4Cl$ in PBS) for 5 min at RT. After this time, EVs-coated coverslips were incubated with blocking solution (5% bovine serum albumin in PBS) for 30 min at RT and then, with either α-PfVps32A, α-PfVps32B or α-PfBro1 polyclonal antibodies labeled with Alexa Fluor 647 or fluorescein isothiocyanate (FITC) (1:100 in all cases) to detect EVs from parasite origin, or anti-GPA (1:100), followed by anti-rabbit Alexa Fluor 488 (1:100) to reveal EVs derived from the RBC plasma membrane. Finally, coverslips were mounted with the appropriate buffer for STORM imaging.

#### Simulations

All simulations were executed on the 'hot' computer cluster of the Max Planck Institute of Colloids and Interfaces. Software used for structure refining and protein docking simulations was the Rosetta software suite version 3.6. Visualizations of protein structures in this paper were produced using pyMol with custom Python scripts. For all data analyses and curvature calculations custom Python scripts were written.

<u>Structure refining</u>

Rough structure predictions of PfBro1 and PfVps32 were acquired from the Phyre2 server[8] (http://www.sbg.bio.ic.ac.uk/phyre2/html/page.cgi?id=index) and subjected to structure refining to prepare both proteins for docking and adapt them to the Rosetta force field. A total of 11,652 PfBro1 and 27,520 PfVps32 structures were generated with a standard FastRelax protocol. Given the rigidity and size of the PfBro1 protein (819 residues), a further refinement in a backrub simulation was performed where 200 structures were generated, of which the best 20 structures from the relax ensemble were chosen based on their total score.

Protein docking

Potential orientations for protein docking of PfBro1 and PfVps32 were previously acquired using the ClusPro web server (https://cluspro.org)[9], omitting the need for Global docking simulations. As input we used two PfVps32 structures in 'open' state and a good scoring PfBro1 structure. For docking simulations these structures were aligned with a prominent prediction from ClusPro using pyMol. To remove any steric hindrances that could be present due to our manual construction of the protein-protein system, a quick restrained relaxation was performed with 96 structures each, the best of which were chosen for the following docking simulations. After performing a prepacking protocol (also with the 96 best scoring structures), local docking simulations were performed generating 6,144 structures for each system. Out of these, the 1,000 lowest scoring structures were filtered using a low-pass filter on an interface score of minus 5 REU. The ca. 20 structures per system that remained after filtering were then further refined by performing additional backrubbing simulations, once again generating 200 structures for each filtered structure.

##### Femtoliter injection

For femtoinjection experiments, GUVs were grown by gel-assisted method from a lipid mixture of POPC, POPS, DSPE-PEG-biotin and DPPE-rhodamine (79:20:1:0.1 mol%). All lipids were obtained from Avanti Polar Lipids (Alabaster, IL, USA). Briefly, a 5% (w/w) polyvinyl alcohol (PVA) solution was prepared in protein buffer (25 mM tris-HCl, pH 7.4, 150 mM NaCl). The PVA solution was spread on a microscope coverslip and then dried for at least 30 min at 50 °C. 10-15 µl of lipids dissolved in chloroform (1 mg/ml) were spread on the dried PVA film and placed under vacuum for 1 h to eliminate the solvent. A chamber was formed with a homemade Teflon® spacer sandwiched between two glasses and filled with protein buffer for 10 min at RT. Then, GUVs were harvested by gentle tapping on the bottom of the chamber and collected using a micropipette without touching the PVA film to avoid sample contamination. To immobilize GUVs, cleaned coverslips were incubated for 20 min at RT with a 1:1 mixture of 1 mg/ml BSA-biotin, 1 mg/ml BSA (both diluted in protein buffer to maintain osmolarity). After incubation, coverslips were washed with distilled water and incubated with 0.005 mg/ml avidin. Subsequently, slides were washed and dried with $N_2$. The functionalized coverslips were maintained in a closed container to avoid binding of dust. For immobilization, the harvested GUVs were deposited in a homemade observation chamber assembled using a Teflon® spacer and let to settle down for at least 10 min.

The micropipettes used to perform the injection were fabricated from thin wall borosilicate capillaries glass with filament (Harvard Apparatus, Holliston, MA, USA) in a pipette puller (Sutter Instruments, Novato, CA, USA) to obtain bee-needle type tips. For the injection experiments, immobilized GUVs were imaged under a Leica TCS SP5 confocal microscope. The micropipette was placed on a mechanical holder attached to a micromanipulator (Sutter Instruments) and then connected to a Femtojet microinjector set (Eppendorf). Injection was performed in a 15° angle, using a pressure of injection of 150 hPa, time of injection of 5.0 s and a compensation pressure of 1 hPa. The solution injected corresponded to a 4× protein mixture stock (2.4 nM PfBro1 and 4.8 nM of either PfVps32A or PfVps32B dissolved in 1× buffer) and 0.03 mg/ml PEG-FITC to monitor injection.

#### IgG purification and RBC pre-depletion

Rabbit polyclonal antibodies against the studied proteins were diluted 2:1 in protein G binding buffer (Pierce, UK) and loaded into protein G Sepharose columns (GE Healthcare, USA). Columns were washed 4 times with binding buffer and IgGs were eluted in 10 ml protein G elution buffer (Pierce, UK). Neutral pH was restored with Trizma pH 9.0. To eliminate RBC-reactive antibodies, samples were incubated for 1 h with RBCs type B+ (100 µl RBCs per ml of IgG purified fraction), then pelleted at 800× g for 10 min and the pellet discarded.

**Generation of *Pfvps60* KO strain**

Homology regions (HR) of the 5'UTR (HR1, spanning positions −762 to −243 from the *Pfvps60* start codon) and 3'UTR (HR2, spanning positions 1,012 to 1,553 from the *Pfvps60* start codon) were PCR amplified using genomic DNA purified from a *P. falciparum* 3D7 culture synchronized at late stages. Primers used for PCR amplification are listed in table S2. The generated HR1 and HR2 were cloned by ligation using restriction sites *SpeI* and *AflII* (HR1), and *EcoRI* and *NcoI* (HR2) into a modified pL6-*egfp* donor plasmid [10] in which the *yfcu* cassette had been removed. The single guide RNA (sgRNA) specific for the *Pfvps60* gene and targeting the sequence near the 5' end (sgRNA 5', position −225, −206) were generated by cloning annealed oligonucleotides into the *BtgZI* site to generate the pL7-*Pfvps60*_KO_sgRNA3' plasmid. On the other hand, the pDC2-Cas9-U6-h*dhfr* vector [11] was modified by cloning a sgRNA specific for the sequence near the 3' end (sgRNA 3', position 980, 999) into the BtgZI site of this plasmid to generate the pDC2-Cas9-U6-h*dhfr*-pfvps60_KO_sgRNA3' plasmid. All guides were cloned using the In-Fusion system (Clontech, Japan).

For transfection of 3D7 rings, 60 µg of circular pDC2-Cas9-U6-h*dhfr*-pfvps60_KO_sgRNA3' plasmid and 30 µg of linearized (with *PvuI*) donor plasmid were precipitated, washed and resuspended in 30 µl of sterile 10 mM Tris, 1 mM EDTA (TE) buffer. Then, plasmids were diluted in 370 µl of Cytomix buffer

24 h after transfection, cultures were selected with 10 nM WR99210 for 4 consecutive days [12]. To validate the integration of the plasmids, a PCR analysis was performed

##### Growth inhibition activity (GIA) assays

Synchronized parasites were cultured as described above. Late-form trophozoite and schizont stages were purified in 70% Percoll and parasites were brought to 4% hematocrit and 1% parasitemia by dilution with fresh RBCs. Samples were prepared by taking RBC-depleted IgG purified fractions and buffer exchanging three times into RMPI using Amicon 10K MWCO centrifugal filtration devices. Cultures were incubated with different concentrations of the samples in a 96-well plate for 48 h under the conditions described previously. For inhibition determination, samples were diluted 1:100 in PBS, nuclei were stained with 0.1 µM Syto11 (ThermoFisher, USA) and read by excitation through a 488 nm laser at 50 mW power and emission collection with a 530/30 bandpass filter with a BD LSRFortessa flow cytometer (Becton, Dickinson and Company, New Jersey, USA). Forward- and side-scatter in a logarithmic scale were used to gate the RBC population.  Acquisition was configured to stop after recording 20,000 events within the RBC population.
