## Supplementary material for "The ESCRT-III machinery participates in the production of extracellular vesicles and protein export during *Plasmodium falciparum* infection": Table S2

| Name | Sequence (5'-3') | Use |
|---|---|---|
| HR1_vps60_S | actagtgttgttctatagtattaattggtacgg | Amplification HR1 Pf*vps60* gene |
| HR1_vps60_AS | cttaagataaatcaactttatgaattaaaaaaaagg | Amplification HR1 Pf*vps60* gene |
| HR2_vps60_S | gaattcctagctttaattgaagatagtatatgg | Amplification HR2 Pf*vps60* gene |
| HR2_vps60_AS | ccatgggtccacattaatttgtttttatttggg | Amplification HR2 Pf*vps60* gene |
| sgRNA5'_vps60_S | taagtatataatatttgagggagaaagtaaacatagttttagagctagaa | Generation of sgRNA5' |
| sgRNA5'_vps60_AS | ttctagctctaaaactatgtttactttctccctcaaatattatatactta | Generation of sgRNA5' |
| sgRNA3'_vps60_S | taagtatataatattctaaatttggatgaacaagagttttagagctagaa | Generation of sgRNA3' |
| sgRNA3'_vps60_AS | ttctagctctaaaactcttgttcatccaaatttagaatattatatactta | Generation of sgRNA3' |
| P1_vps60 | tcaagggtaagaaagataaaaccattggg | PCR diagnostics KO *Pfvps60* |
| P2_vps60 | tgaggatgataaaatatttccaatttcgg | PCR diagnostics KO *Pfvps61* |
| P3_vps60 | gtgccttttctggtaaagggttgg | PCR diagnostics insertion donor plasmid |
| P4_vps60 | agggcatatgtaaatggccaaagg | PCR diagnostics insertion donor plasmid |
